# Bactofilins form non-polar filaments that bind to membranes directly

**DOI:** 10.1101/617639

**Authors:** Xian Deng, Andres Gonzalez Llamazares, James Wagstaff, Victoria L. Hale, Giuseppe Cannone, Stephen H. McLaughlin, Danguole Kureisaite-Ciziene, Jan Löwe

**Author notes:** authors contributed equally (Xian Deng, Andres Gonzalez Llamazares). corresponding author: Jan Löwe, MRC Laboratory of Molecular Biology, Francis Crick Avenue, Cambridge CB2 0QH, UK.

## Abstract

Bactofilins are small beta-helical proteins that form cytoskeletal filaments in a range of bacteria. Bactofilins have diverse functions: filaments in *Caulobacter crescentus* are involved in cell stalk formation whereas *Myxococcus xanthus* filaments aid chromosome segregation and motility. The precise molecular architecture of bactofilin filaments has remained unclear. Here we revealed by sequence analyses and electron microscopy that in addition to wide distribution across bacteria and archaea, bactofilins are also present in a few eukaryotic cells such as oomycetes. The sole bactofilin from *Thermus thermophilus* was demonstrated to form constitutive filaments and cryo-EM analysis revealed that protofilaments formed through end-to-end association of the beta-helical domains. Using a nanobody against *Thermus* bactofilin we determined the near-atomic filament structure, showing that the filaments are non-polar, with subunits arranged head-to-head and tail-to-tail. A polymerisation-impaired mutant F105R, that disrupts one of the two protofilament interfaces, enabled crystallisation. The crystal structure also revealed non-polar protofilaments, and the dominance of the beta-stacking interface that formed despite the inhibiting mutation. To confirm the generality of the lack of polarity, we performed co-evolutionary analysis of a large set of sequences. Finally, using *Thermus* bactofilin, we determined that the N-terminal disordered tail of the protein is responsible for direct binding to lipid membranes both on liposomes and by electron cryotomography in *E. coli* cells. The tail is conserved, suggesting that membrane binding is likely a general feature of these very common but only recently discovered filaments of the prokaryotic cytoskeleton.

## Introduction

Most bacteria and archaea contain protein filaments that provide functionality on length scales beyond that of their subunits and that have been collectively called prokaryotic cytoskeletons (Wagstaff and Löwe, 2018). Amongst those, cytomotive filaments of the actin and tubulin types have attracted most attention as they engage in conserved cell biological functions such as cell division, DNA segregation and cell shape determination (Amos and Löwe, 2017). Cytomotive filaments are characterised by complex dynamics driven by cycles of coupled polymerisation and depolymerisation, nucleotide hydrolysis and exchange events, and conformational changes within each monomer (Michie and Löwe, 2006; Miraldi et al., 2008; Wagstaff et al., 2017).

Most prokaryotes also encode protein filaments that do not belong to the actin and tubulin families and that do not function through filament dynamics enabled by filament-bound nucleotide hydrolysis, instead acting as molecular scaffolding (Lin and Thanbichler, 2013). Martin Thanbichler and co-workers recognised in 2010, after several earlier sightings, the existence of a conserved and widespread family of bacterial proteins that form constitutive and very stable filaments *in vitro* and which perform scaffolding roles in cells, they named these proteins “bactofilins” (Kühn et al., 2010). Bactofilins always contain a small domain of about 110 amino acids (PFAM domain PF04519 “bactofilin”, originally annotated DUF 583, pfam.xfam.org), which has been recognised as being present in many bacterial genomes, often within multiple genes (Lin and Thanbichler, 2013). Most bactofilin sequences consist of one bactofilin domain flanked by presumably disordered proline-rich tails at the N- and C-terminal termini, although this is not universal: enterobacterial bactofilins, including *Proteus Mirabilis* CcmA, have predicted transmembrane helices in the N-terminal tail (Hay et al., 1999; Kühn et al., 2010). Bactofilin filaments are very stable, being largely insensitive to pH, salt concentration and chelating agents and hence are always filamentous when purified from source or expressed heterologously (Koch et al., 2011; Kühn et al., 2010).

The structure of the conserved bactofilin domain was solved by the first ever *de novo* solid state NMR (ssNMR) atomic structure determination (Shi et al., 2015), with supporting evidence coming from sequence-based modelling and electron microscopy of the filaments (Kassem et al., 2016; Vasa et al., 2015; Zuckerman et al., 2015). The bactofilin domain has a right-handed beta-helical fold, with 6 windings of ~17 amino acid residues producing triangular-shaped monomers that measure roughly 3 nm along the beta-helical axis. So far, it has not proved possible to determine the filament structure of bactofilins.

Understanding of bactofilin function is so far limited to a small number of examples. *Caulobacter crescentus* bactofilins BacA and BacB were identified in a localisation screen for proteins involved in stalk formation in these asymmetric cells (Kühn et al., 2010). BacAB are expressed throughout the cell cycle and condense at the stalk-forming site within the cell at the onset of S phase. These bactofilins have been found to directly interact with the cell wall synthesis enzyme PbpC, and deletion of either the filaments or PbpC produces much shorter stalks. In the cells, BacA and BacB have been reported to form filaments or sheets close to the cell’s inner membrane and over-expression deforms cells, making it possible that the filaments have intrinsic curvature and bind membranes. Some biochemical evidence suggested that BacAB are peripheral membrane proteins (Kühn et al., 2010).

Other well-investigated bactofilins are BacM, N, O and P from *Myxococcus xanthus*. In contrast to *Caulobacter*, where BacA and BacB seem to have overlapping functions and copolymerise, *Myxococcus* bactofilins have been reported to have at least three different functions. A *bacM* gene knock-out showed ‘crooked’ cells that have increased sensitivity to antibiotics (Koch et al., 2011). BacNOP, in contrast, co-polymerise into filaments that constrict ParABS chromosome segregation proteins to subpolar regions of the cells (Lin et al., 2017). BacP has also been reported to be involved in type IV pilus localisation together with the GTPase SofG, both being important for the direction of motility of *Myxococcus* cells (Bulyha et al., 2013).

*Helicobacter pylori* contains a single bactofilin called CcmA. A *ccmA* gene knock-out completely abolished the characteristic helical shape of the cells and a model has been put forward in which CcmA bactofilin filaments position lytic endopeptidases Csd1-3 in the periplasm, to remodel the shape-giving cell wall (Blair et al., 2018; Sycuro et al., 2010). Bactofilins have also been described in another spirochete, *Leptospira biflexa*, with one type of bactofilin being seemingly responsible for specific helical pitch parameters of the cells (Jackson et al., 2018).

The first sighting of a bactofilin, although not described as such then, was CcmA from *Proteus mirabilis*, where it is involved in cellular motility (Hay et al., 1999), and a similar functional context was described for CcmA / bactofilin proteins in *Vibrio parahaemolyticus* (Gode-Potratz et al., 2011) and *Bacillus subtilis* (El Andari et al., 2015; Rajagopala et al., 2007).

Clearly, more, and more precise, functional and cell biological investigations will be needed and we hope to be able to facilitate this here with our work determining the atomic structure of the bactofilin filament from *Thermus thermophilus by* cryo-EM. The filament is composed of domains stacked so that a continuous beta-helical filament results. The subunits are arranged head-to-head, resulting in a filament that has no polarity. This finding was confirmed by crystallography and co-evolutionary analysis. We show that the filaments bind directly to membranes *in vitro* and when heterologously expressed in *E. coli* cells, and that this interaction is enabled by a short conserved and hydrophobic motif in the N-terminal tail. Finally, we also show that polymerising bactofilins are not restricted to bacteria, with the visualisation of bactofilin filaments from *Phytopthora infestans*, an oomycete eukaryote.

## Results

### Bactofilin domains are found across the tree of life

The presence of bactofilins has been reported and experimentally verified in several bacterial clades (Figures 1A and B). In order to assess further the distribution of the conserved bactofilin domain we searched for the presence of the conserved PF04519 / DUF 583 domain in a curated set of prokaryotic genomes (Mendler et al.), annotated with a standardised phylogenomic taxonomy (GTDB v86) (Parks et al., 2018) (resulting tree: Supplementary Data D1). Using a standardised taxonomy allowed for a meaningful assessment of the distribution of individual genes across clades, because clades of equal taxonomic rank represent roughly comparable levels of genomic divergence. We found that bactofilin domains were common, defined as being present in more than 20% of genomes, in 82 of the 114 phylum-level clades in the standardised taxonomy (Figure 1A). We conclude that bactofilins are very widely distributed within bacteria. We also investigated the presence of bactofilin domains in the domain Archaea in the same way, finding that many archaeal genomes harbour bactofilin domains. Within the phylum-level clade Halobacterota more than 80% of genomes contained at least one bactofilin domain hit. It remains to be experimentally verified that these archaeal sequences encode polymerising bactofilin.

**Figure 1.**
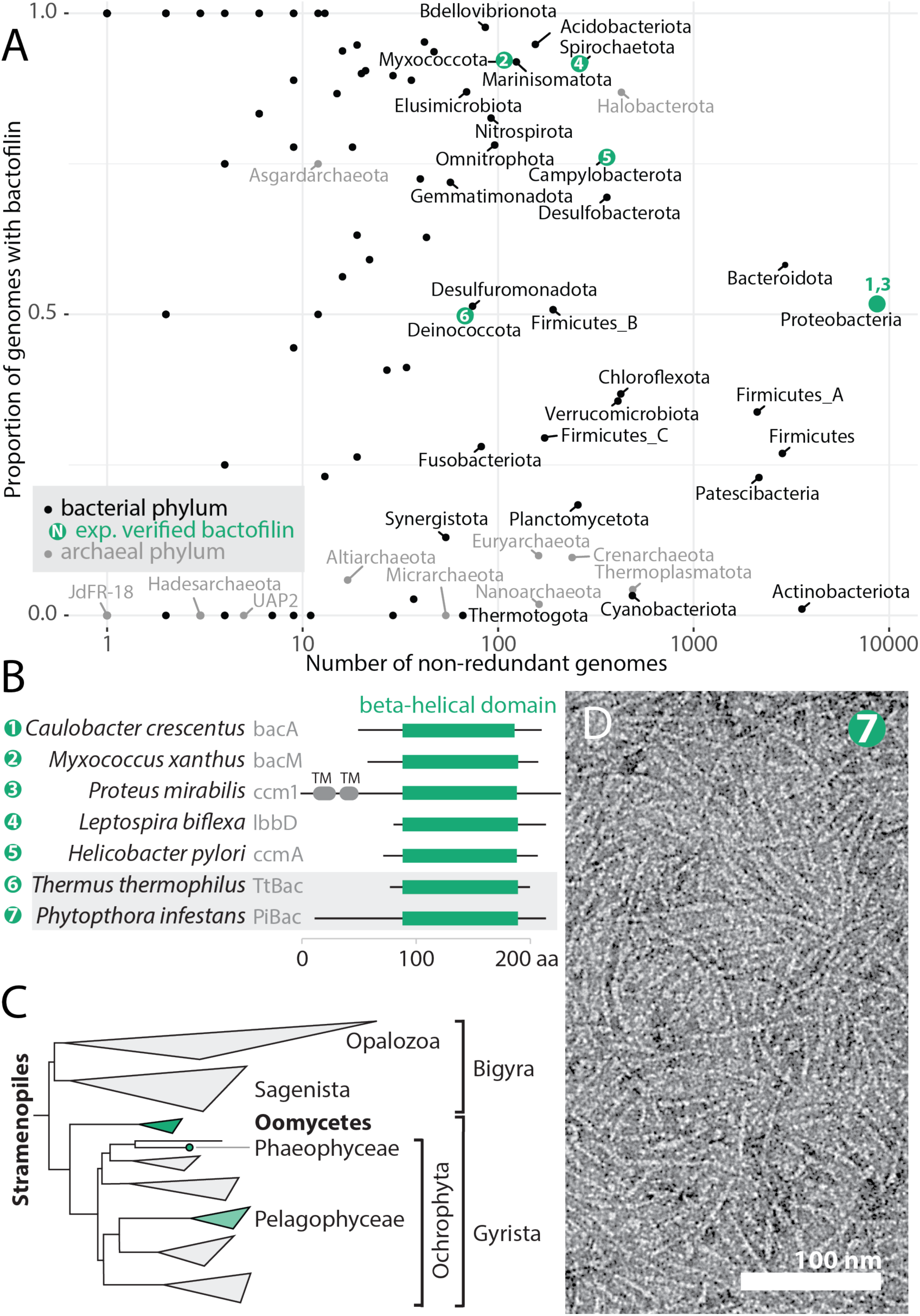
Bactofilins are highly conserved and widespread across bacteria and archaea, and also occur in some eukaryotic organisms. A) Phylum level clades from the GTDB taxonomy v86 (Parks et al., 2018) of bacteria (black) and archaea (grey) are plotted as points with the number of non-redundant genomes in the clade against the proportion of genomes with at least one bactofilin hit. Bactofilin hits were HMMSEARCH results with PFAM PF04519, using an E-value cutoff of 1e-10 for bacteria, and 1e-6 for archaea, chosen so as to retain clusters of similar, convincing, sequences in each case. Pre-computed HMMSEARCH results were retrieved from the AnnoTree server (Mendler et al.). Clades containing the validated bactofilins in (B) are shown in green, and numbered to correspond to (B). B) Domain schematic of selected experimentally investigated bactofilins. Beta helical domain (green) is shown as regions aligning to the beta helical regions in *C. crescentus* BacA (Shi et al., 2015) and *T. thermophilus* TtBac structures. UniProt accessions for proteins shown, 1: Q9A753, 2: Q1CVJ5, 3: B4F0H9, 4: B0SPX3, 5: corresponds to GenBank CP001173.1 1607606:1607196 (Uniprot shows incorrect start codon see (Sycuro et al., 2010), 6: Q72HS6, 7: D0N980. C) A phylogenomic tree of the eukaryotic Stramenopile clade (Derelle et al., 2016). The subclades with genomes containing putative bactofilins are highlighted in green, the lighter shade of the Pelagophycaeae corresponds to a patchier distribution, see text. D) Negative stain electron micrograph showing filaments formed by recombinant bactofilin PiBac (PITG_07992) from the oomycete *Phytopthora infestans*.

While browsing the PFAM entry for the bactofilin domain PF04519 we noticed that several hits are listed within the domain Eukarya (El-Gebali et al., 2019). We confirmed that the sequences looked like *bona fide* bactofilins and undertook a more thorough search of eukaryotic genomes for bactofilin domains. We found convincing bactofilin-like sequences in two taxonomic clusters, one within Stramenopiles (a deeply rooted eukaryotic clade, Figure 1C) and another within the Ascomycete fungi (Supplementary Data D2). We recombinantly expressed a putative bactofilin gene (Uniprot D0N980, PITG_07992, herein “PiBac”) from the economically important Stramenopile plant pathogen *Phytopthora infestans*, a member of the Oomycete group, and found that it indeed forms bactofilin-like filaments (Figure 1D). Published data show that the mRNA encoding PiBac is expressed before and during spore formation (Supplementary Figure S1), and that the gene is conserved in context across the spore-forming Oomycetes (Supplementary Figure S2) (Ah-Fong et al., 2017). Therefore, bactofilin domains retaining the ability to polymerise are conserved in eukaryotic genomes, are expressed, and likely play functional roles. The widespread distribution of polymerising bactofilin domains underscored the need for a fuller structural understanding of bactofilin polymers.

### Bactofilin from Thermus thermophilus (TtBac) forms completely beta-helical filaments

Bactofilin from *Thermus thermophilus* presented itself as a promising candidate for structural determination due to its short N- and C-terminal tails flanking the conserved bactofilin domain and the biological tractability of the source organism. Purification of recombinantly expressed His_6_-tagged TtBac (H_6_-TtBac, Supplementary Table T2) under denaturing conditions and subsequent refolding allowed visualisation of TtBac filaments by negative stain electron microscopy and confirmed the *in silico* work that predicted the presence of a putative polymerising bactofilin in *Thermus thermophilus* (Figure 2A). Additionally, the H_6_-TtBac filaments recapitulated previously reported 2D sheets that seem to be an intrinsic consequence of fibrillar bactofilin assemblies.

**Figure 2.**
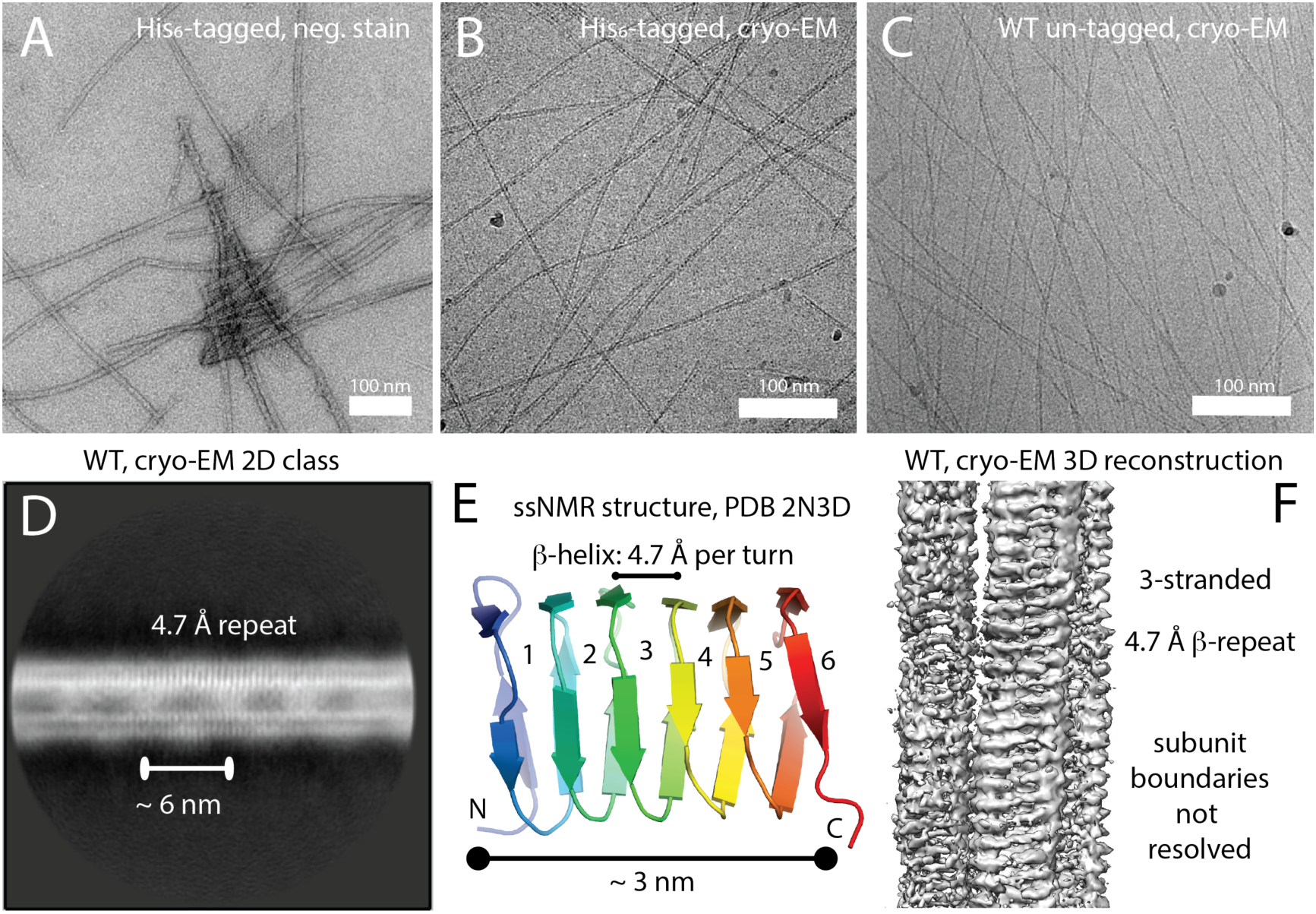
Bactofilin from *Thermus thermophilus* forms filaments that show a continuous beta-stacking repeat of 4.7 Å. **A)** Hexahistidine-tagged and refolded bactofilin from *Thermus thermophilus* (H_6_-TtBac) forms filaments and 2D sheets as determined by negative stain electron microscopy (EM). B) The same protein shown using electron cryomicroscopy (cryo-EM). Note differing filament thicknesses caused by varying protofilament numbers. C) Untagged TtBac-WT protein expressed in *E. coli* and purified using a centrifugation protocol, imaged by cryo-EM, also showing protofilament numbers of 2-4. D) 2D class average calculated using RELION (Scheres, 2012), showing a continuous repeat of 4.7 Å perpendicular to the filament axis. Note the upper strand shows stronger density, most likely because many filaments averaged have 3 protofilaments. Also note that there are fuzzy bridges visible between the protofilaments, roughly 6 nm apart. E) Ribbon representation of the *Caulobacter cresentus* BacA bactofilin structure determined by solid state NMR (ssNMR) (Shi et al., 2015). Taking D) into account, the structure suggests that bactofilin protofilaments are made by stacking the beta-helical domains head-to-head and tail-to-tail. The 6 nm fuzzy bridges already suggest that the structure might not be polar as it is difficult to produce those repeats lengths from a 3 nm long beta-helical domain otherwise. F) Preliminary RELION helical 3D reconstruction using the cryo-EM data (He and Scheres, 2017), revealing the structure of a 3-stranded TtBac filament. Because the individual domains show no features indicating their ends, the alignment does not converge and resolution remains low because the subunits cannot be registered correctly along the filament axis.

In order to isolate full length, unmodified, TtBac (TtBac-WT) we developed a protocol based on a series of gradient centrifugation steps adapted from previous work on *Myxococcus xanthus* BacM (Koch et al., 2011). Visualisation by cryo-EM of both H_6_-TtBac and natively purified TtBac-WT filaments showed that the fibrils were prone to bundling and persisted unchanged over very wide ranges of pH and salt concentrations. Preparing the filaments at pH 11 allowed production of cryo-EM grids with largely un-bundled filaments that were amenable to structure determination but otherwise looked like filaments at more physiological pH values (Figures 2B & C).

Many 2D class averages obtained with RELION of TtBac-WT showed two lines or protofilaments of helical fibrils with twice as much signal in the top protofilament, suggesting that the majority of assemblies present in the grids were in fact made out of three protofilaments (Figure 2D). The class averages all showed strong vertical striations within the protofilaments, spaced 4.7 Å apart, the distance between beta strands in a sheet. This indicated that the bactofilin subunits arranged into continuous beta-helical protofilaments, as suggested previously based on low resolution work and modelling (Vasa et al., 2015; Zuckerman et al., 2015). Surprisingly, the filaments also showed a weak ~ 6 nm repetitive feature visible between and in the protofilaments (Figure 2D), roughly twice the size of a BacA monomer as previously determined by ssNMR (Shi et al., 2015) (Figure 2E). This indicated that the repeating unit of the filament might be a head-to-head dimer making polarity within the filament impossible.

Subsequent 3D helical reconstruction of TtBac-WT filaments with RELION (He and Scheres, 2017) yielded a preliminary low-resolution structure that confirmed both the 3-strandedness and the β-helical nature of the subunits, but did not allow any atomistic description of the monomer (Figure 2F). The inability to reach higher resolution was mainly caused by the similarity between each β-helical winding, which impeded the correct positioning of the subunits within each protofilament. Therefore, the relative positions of subunits remained unclear and the issue of polarity or lack thereof could not be conclusively resolved.

### A near atomic cryo-EM structure of the TtBac bactofilin filament bound to a nanobody shows it to be non-polar

In an attempt to provide a solution to the computational problem of the subunit register along each protofilament, we raised *Lama glama* nanobodies against TtBac-WT. After selection of the suitable antibody clone NB4, cryo-EM of the wildtype filaments incubated with the anti-TtBac-WT nanobody showed fully decorated fibrils that formed superhelical structures unfit for structure determination (Figure 3A). Therefore, by interpreting the conserved structure of nanobodies, we designed 4 mutations at the opposite end from the CDR loops of the nanobody to counteract the bundling —L13S, Q15D, K45D, and K66D on NB4-mut2 — which almost completely abolished superhelicity (Figures 3B, C & D). Images were clear enough to directly derive approximate helical parameters for image reconstruction (Figure 3D). 2D class averages of the decorated filaments showed almost exclusively filaments with two protofilaments, clear signal for each nanobody at a spacing of 5.7 nm and the two beta-helical protofilaments (Figure 3E).

**Figure 3.**
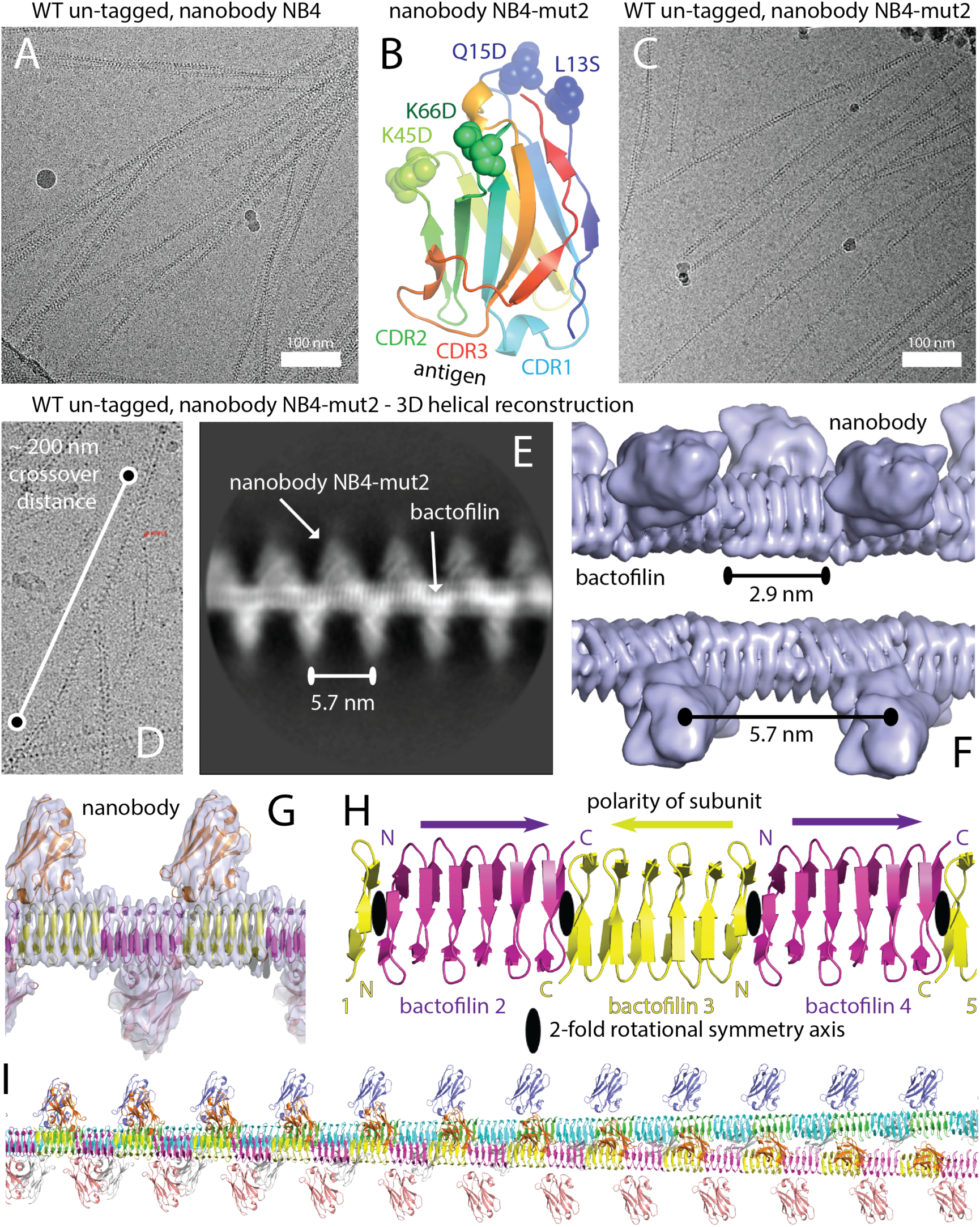
Cryo-EM structure of bactofilin from *Thermus thermophilus* (TtBac) shows a head-to-head non-polar filament. A) In order to overcome the register problems highlighted in Figure 2D-E, a nanobody against TtBac was raised, purified and added to the filaments before cryo-EM imaging. The nanobody clearly binds but causes severe bundling. B) Assuming that bundling is caused by the antigen-distal surface of the nanobody, four mutations were introduced, yielding NB4-mut2. C) Using the modified nanobody NB4-mut2 bundling is significantly reduced and cryo-EM images show many single filaments. D) Estimation of the crossover distance (corresponding to a 180° twist of the filament) to over 2000 Å, indicating only very slowly twisting filaments of about 3-4 ° per 6 nm helical rise. E) Reference-free 2D class calculated using RELION, showing the altered appearance of the TtBac bactofilin filaments after addition of the NB4-mut2 nanobody (compare to Figure 2D). The nanobody densities appear at a repeat distance of 5.7 nm, clearly indicating a non-polar arrangement of the subunits in each protofilament. F) Preliminary low-resolution (5-6 Å) 3D reconstruction using RELION, showing clearly the continuous beta-helical TtBac bactofilin filament pair and the nanobodies attached at 5.7 nm distances to each other. G) Final density with fitted (but not refined) atomic models at around 4 Å resolution. Correctness of the handedness of the structure was verified using the crystal structure fit of the nanobody atomic model. H) Ribbon representation of the arrangement of individual TtBac subunits in each protofilament. Subunits are arranged in a head-to-head-to-tail manner, with N- and C-termini coming together at alternating interfaces (N-C and C-C, leading to the 5.7 nm repeat of the nanobody binding). The two-fold axes at the N-N and C-C interfaces are nearly aligned, leading to a slowly twisting, overall helical filament. I) Atomic model of a longer stretch of the double-helical filament. The entire double filament has additional 2-fold symmetry axes going between the two protofilaments.

Thus —with the aid of nanobody NB4-mut2— by averaging ~ 346,000 helical segments and applying and refining the final helical parameters in RELION helical image reconstruction (He and Scheres, 2017), the structure of TtBac-WT:NB4-mut2 filaments was solved to a nominal resolution of 3.4 Å (Figures 3F & G, Supplementary Table T1, Supplementary Figure S3, 4.2 Å against atomic model). Fitting the previously determined structure of bactofilin BacA from *Caulobacter* (Shi et al., 2015), revealed close similarity between BacA and TtBac-WT (Figure 3G). As in BacA, TtBac is made up of 6 right-handed windings of triangular parallel β-sheet structures with an exclusively hydrophobic core —a main contributor to the extreme stability of bactofilins— and two disordered terminal tails since the ordered part of TtBac only extends from amino acids 12 to 112. Similar to the highly conserved glutamine and asparagine hydrogen bonds that give an increased sturdiness to cross-β amyloid fibres, bactofilins contain a bonding network comprised of glutamate and aspartate placed diametrically above or below lysine and arginine. The structural simplicity of the TtBac filaments means that amino acid composition alone reflects their three main characteristics: an hydrophobic core consisting of leucine, valine, or alanine (34 % of residues); a strong interaction network between surface glutamates, arginines and lysines, together comprising 28 % of amino acids; and a triangular arrangement of β-sheets with glycines (13 %) at each sharp corner.

Primarily, the cryo-EM structure provides the first insight into the filamentous arrangement of bactofilin, showing that the monomers are ordered in a non-polar fashion with alternating head-to-head (N-terminal to N-terminal, N-N) and tail-to-tail (C-terminal to C-terminal, C-C) interfaces (Figure 3H). This in turn produces 2-fold symmetry at each homoterminal N-N and C-C interface. The interfaces themselves have a very small misalignment of their 2-fold axes, translating into a slight helical twist of 4.89° per dimer (Figure 3I, Supplementary Figure S4).

### The crystal structure of a TtBac polymerisation-impaired mutant confirms the non-polar filament architecture

Although the filamentous structure converged to near-atomic resolution, the presence of anisotropy in the cryo-EM map, caused by the strong 4.7 Å repeat in a single direction made it difficult to confidently assert the exact conformation of amino acids and other characteristics of the structure. Therefore, in order to validate the cryo-EM reconstruction, we aimed to solve the structure of non-polymerising TtBac versions through X-ray crystallography. By introducing the mutation F105R (Zuckerman et al., 2015) into the C-terminal interface of TtBac we aimed to obstruct polymerisation at this interface while preserving the head-to-head N-N contacts and thus producing a dimer of TtBac(F105R) (Figure 4A). To further aid purification and crystallisation we added a C-terminal His_6_-tag and removed the first 10 N-terminal residues, which, according to the unresolved density in the 2D classes, seemed to be implicated in protofilaments binding to each other (Figure 3F). Indeed, the easy handling of ΔN-TtBac(F105R) was in stark contrast to the difficult purification of TtBac-WT. Moreover, visualisation of the polymerisation-impaired and non-bundling mutant by negative stain EM showed a lack of filaments when tested at fairly high concentrations (~2 mg/ml; not shown).

**Figure 4.**
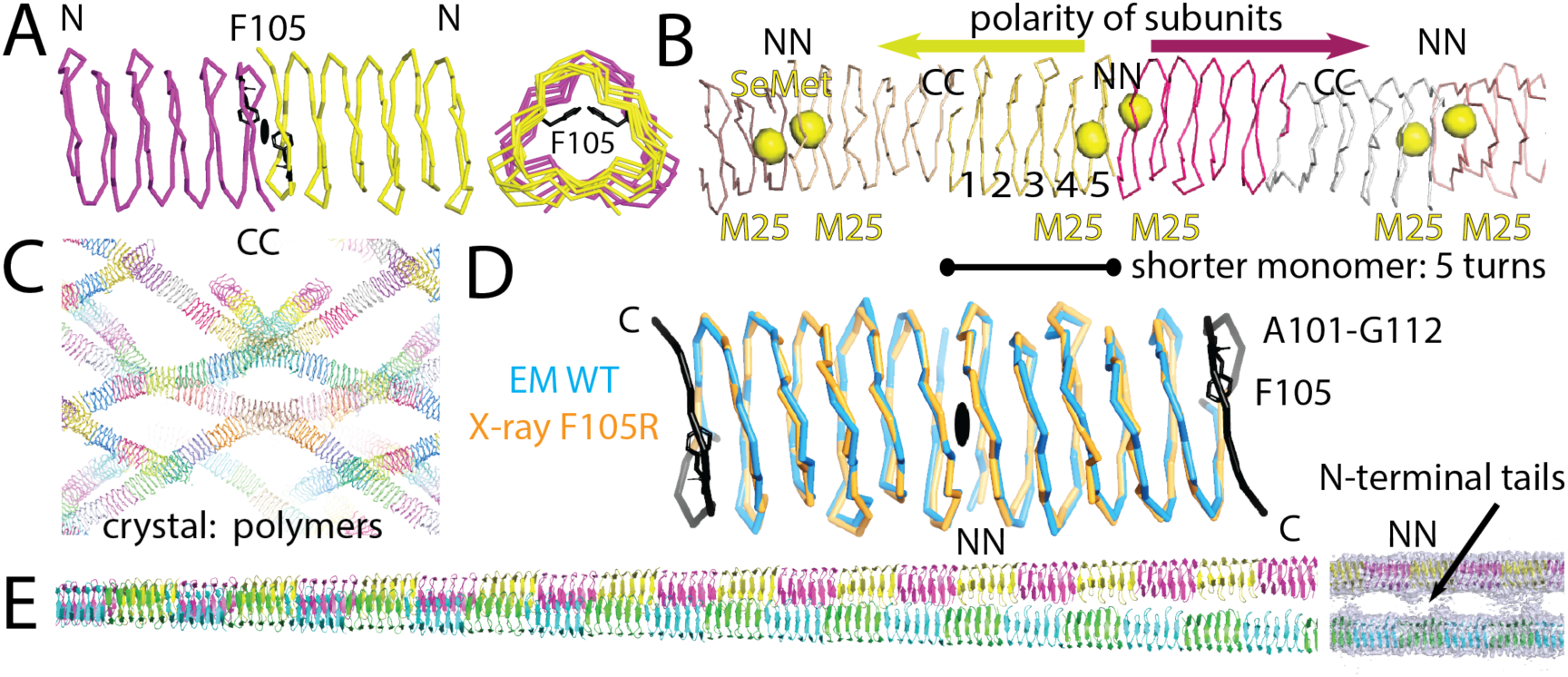
A polymerisation-impaired TtBac mutant crystallises as non-polar filaments with reduced subunit length. A) Based on previous studies (Zuckerman et al., 2015), F105 in TtBac was mutated to R in order to produce a version that does not polymerise under normal conditions. Normally, F105 interacts with itself across the C-C interface along the TtBac protofilament and changing this residue to charged arginine impairs normal filament formation. TtBac(F105R) can be purified easily, especially when the disordered N-terminal region is also removed. B) Crystal structure of ΔN-TtBac(F105R). The structure was solved by SeMet SAD phasing and the resulting signals of the single Met25 residue all lie very close to each other signalling the N-N interface and indicating non-polar head-to-head-to tail arrangement. C) The very large unit cell of the TtBac crystals contains several strands of filaments, formed from 32 molecules per asymmetric unit. D) The mutant crystallised as a distorted filament because the F105R mutation caused residues 101-112 to become disordered, allowing an entirely new C-C beta-helical interface to form. This re-arrangement indicates the power of beta-stacking interfaces as they are also occurring in amyloid structures, for example and that lead to the extraordinary stability of bactofilin filaments. E) Final atomic model of the double-helical TtBac bactofilin filament. Using subtraction in RELION (Bai et al., 2015), the nanobody density was removed and the atomic model refined against the crystal structure was used as guidance to produce a reliable atomic model that was real-space refined against the cryo-EM density. Note the position of the inter-protofilament density bridges, occurring at the locations of N-N interfaces, indicating that it is the N-terminal tails of TtBac that hold the two protofilaments together in double helical filaments.

The crystal structure of ΔN-TtBac(F105R) was solved by making use of the single N-terminal Met25 for selenomethionine (SeMet) SAD phasing (Figure 4B, Supplementary Table T1). In the solved structure, the un-mutated N-N interface was intact and head-to-head, as unequivocally shown by the positions of the anomalous selenium peaks. The map also showed that, although at 2 mg/ml bactofilin filaments were not present, crystallisation occurred through formation of a C-C interface, leading to filaments in the crystals, with 32 bactofilin subunits per asymmetric unit of the crystals (Figure 4C). This unexpected polymer was formed by unwinding of the sixth β-helical turn, essentially flipping out the R105 residue that would otherwise have impaired polymerisation (Figure 4D). This not only suggests that F105R did indeed impair normal polymerisation but also serves to show the outstanding propensity of β-stacks to form, as for example in amyloid fibres.

Aside from confirming the lack of polarity within the protofilaments, the structure of ΔN-TtBac(F105R) also suggested that the formation of higher-order structures such as double filaments may well be caused by the N-terminal residues 1-10. The large unit cell of the crystals contains well-dispersed filaments that do not form any type of doublets or bundles (Figure 4C).

The refined crystallographic model (Supplementary Table T1) was fitted into a new cryo-EM map generated by signal subtraction of the nanobody density (Bai et al., 2015). We added the remaining residues and refined in reciprocal and real-space against the cryo-EM map to obtain a complete and reliable atomic model of the TtBac-WT bactofilin double filament (Figure 4E, Supplementary Table T1). The resulting map again highlighted that the bridges between the protofilaments that make the double filaments are formed by N-terminal residues (Figure 4E, right).

### All bactofilins form non-polar filaments as demonstrated by evolutionary coupling analysis

The high level of conservation between the monomer structures of TtBac and *Caulobacter* BacA (Shi et al., 2015) (PDB ID 3N3D, RMSD 1.7 Å over 96 residues, sequence identity 35 %) suggested to us that the non-polar architecture seen for TtBac filaments may also be widely conserved. To investigate, we aligned 12,646 putative bactofilin beta-helical domain sequences and derived an evolutionary coupling score for each pair of residues in the alignment (Ekeberg et al., 2013). Visualisation of these scores as a heat map produced an overall view of co-evolution within the bactofilin sequence and was compared to calculated C*α* distances in the TtBac monomer structure (Figures 5A & B). As shown by the second diagonal in Figure 5B, there is a high level of co-evolution, and high coupling scores, between amino acids ~17 residues apart, i.e. one turn of the beta-helix, and a weaker line ~34 residues apart (two turns) is also visible. In other words, the β-helical structure of bactofilins, which enforces interactions between the residues exactly above and below, is conserved amongst a very large proportion of the aligned sequences.

**Figure 5.**
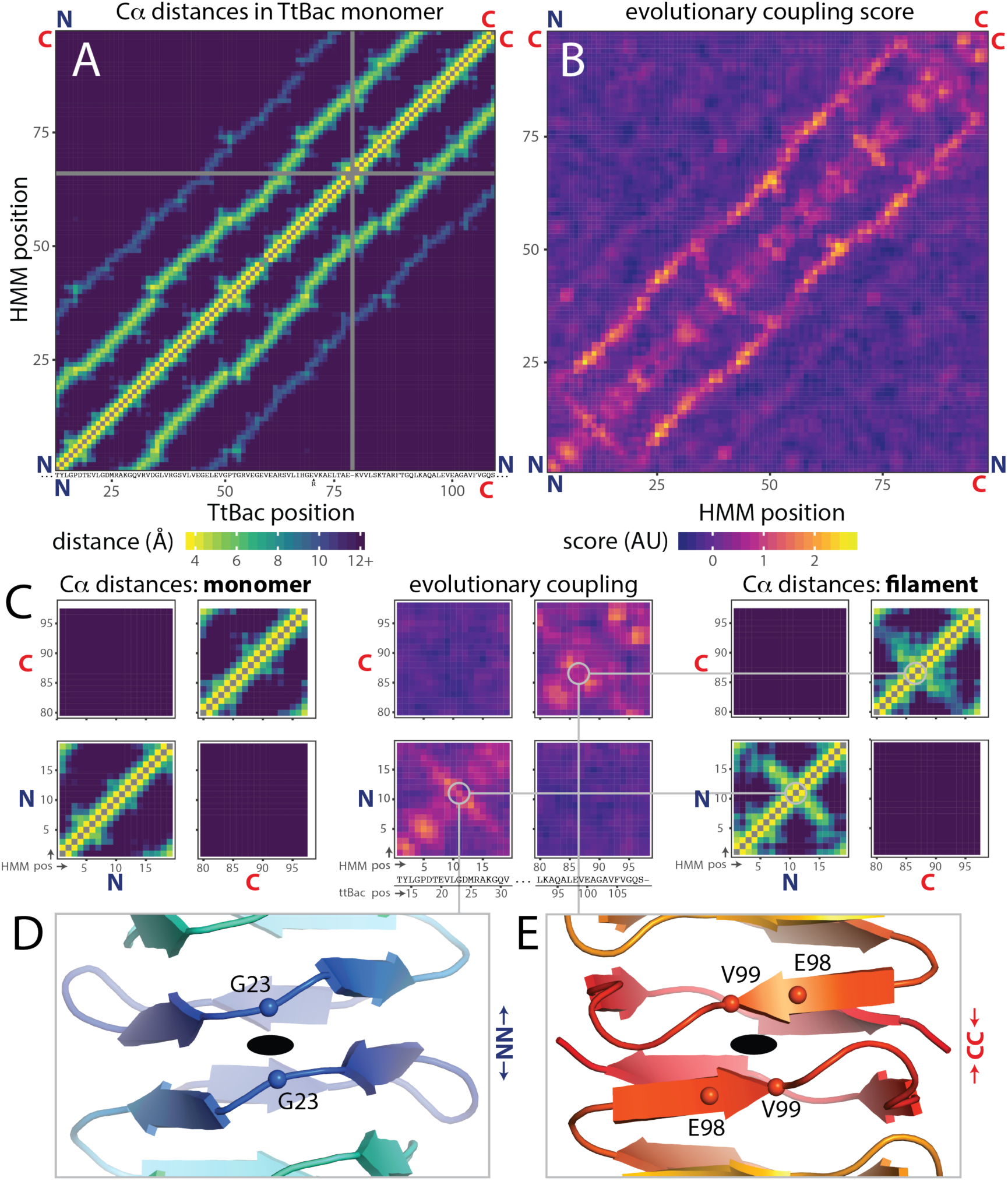
Evolutionary sequence coupling analysis reveals conserved non-polar architecture of bactofilin filaments. A) C alpha distance matrix for TtBac monomer structure within cryo-EM filament structure. The ~17 amino acid repeat becomes obvious. The TtBac sequence was aligned to PFAM HMM PF04519 using HMMALIGN, X-axis numbers are for TtBac sequence, insertions and gaps relative to HMM are indicated below axis, Y-axis is numbered in HMM coordinates. As are all other axes in the figure, except where noted. B) Evolutionary coupling scores calculated for an alignment of 12,646 bactofilin sequences. Scores were smoothed using a 3×3 Gaussian kernel. ~17 amino acid repeat is clear. C) Zoomed plots showing N-N, C-C and N-C interaction regions in distance matrix plot and evolutionary coupling matrix. (Left) zoom in to corners of (A). (Middle) zoom in to (B) with TtBac sequence added beneath. (Right) zoom as for A, but minimum distances between C alphas in filament model are plotted instead. Inter-subunit interactions appear in both the filament distance matrix and coupling matrix as diagonals going through the residues closest to the C2 axis. D) View along C2 axis at N-N inter-subunit interface of TtBac filament model. Chains are displayed in cartoon representation, coloured blue-red from N terminus to C terminus. C2 axis is shown as black oval. C alphas for residues closest to axes are shown as spheres and labelled. Grey lines connect panel to the relevant positions in C. E) As (D) for C2 axis at C-C inter-subunit interface.

As the strength of evolutionary coupling between amino acids is largely dictated by their physical distance, it is reassuring that the heat map of Cα distances for residues in the TtBac monomer in Figure 5A almost perfectly recapitulates the coupling scores observed in Figure 5B. However, the distances between TtBac residues in the monomer do not explain the co-evolution between homo-terminal amino acids observed in the lower left and upper right corners of the coupling heat map (Figures 5B and blown up in Figure 5C, middle). These scores can be explained, though, by taking into account the architecture of the TtBac filament as determined here (Figure 5C, right). In the TtBac filament it is precisely those residues with surprisingly high coupling scores that are brought closer together (than in the monomeric structure) within the head-to-head and tail-to-tail polymer architecture. There is no sign of co-evolution corresponding to interaction between the C- and N-termini (top left/bottom right corners) as would be expected for a polar head-to-tail arrangement.

As the strong coupling scores exclusively indicate interaction between residues within each terminus (N-N and C-C, not N-C) it becomes clear that the non-polar arrangement of the TtBac filament must be highly conserved within bactofilins. The generality of the TtBac filament architecture extends to fine details such as minimal protofilament twist as the position of the orthogonal diagonal in the coupling heat map describing the homo-terminal interaction exactly matches the location of the two-fold symmetry seen in the cryo-EM structure (Figure 5D & E).

### TtBac bactofilin binds to membranes in vitro and in vivo, an activity facilitated by its N-terminal tail

It has previously been shown that bactofilins are located close to membranes in cells *in vivo* (Kühn et al., 2010). Other filaments of prokaryotic cytoskeletons have been shown to directly interact with membranes, either through amphipathic helices or other small membrane targeting signals (MTS) (Pichoff and Lutkenhaus, 2005; Salje et al., 2011; Szeto et al., 2002). Because the N-terminal tail of TtBac was shown here to mediate interactions between the two protofilaments (Figure 4E), while being disordered, we hypothesised that it might contain hydrophobicity that is normally used to interact directly with membranes. Indeed, when aligning a subset of well-characterised homologous bactofilin sequences, a conserved hydrophobic motif emerged at the very N-terminus of these sequences (Figure 6A).

**Figure 6.**
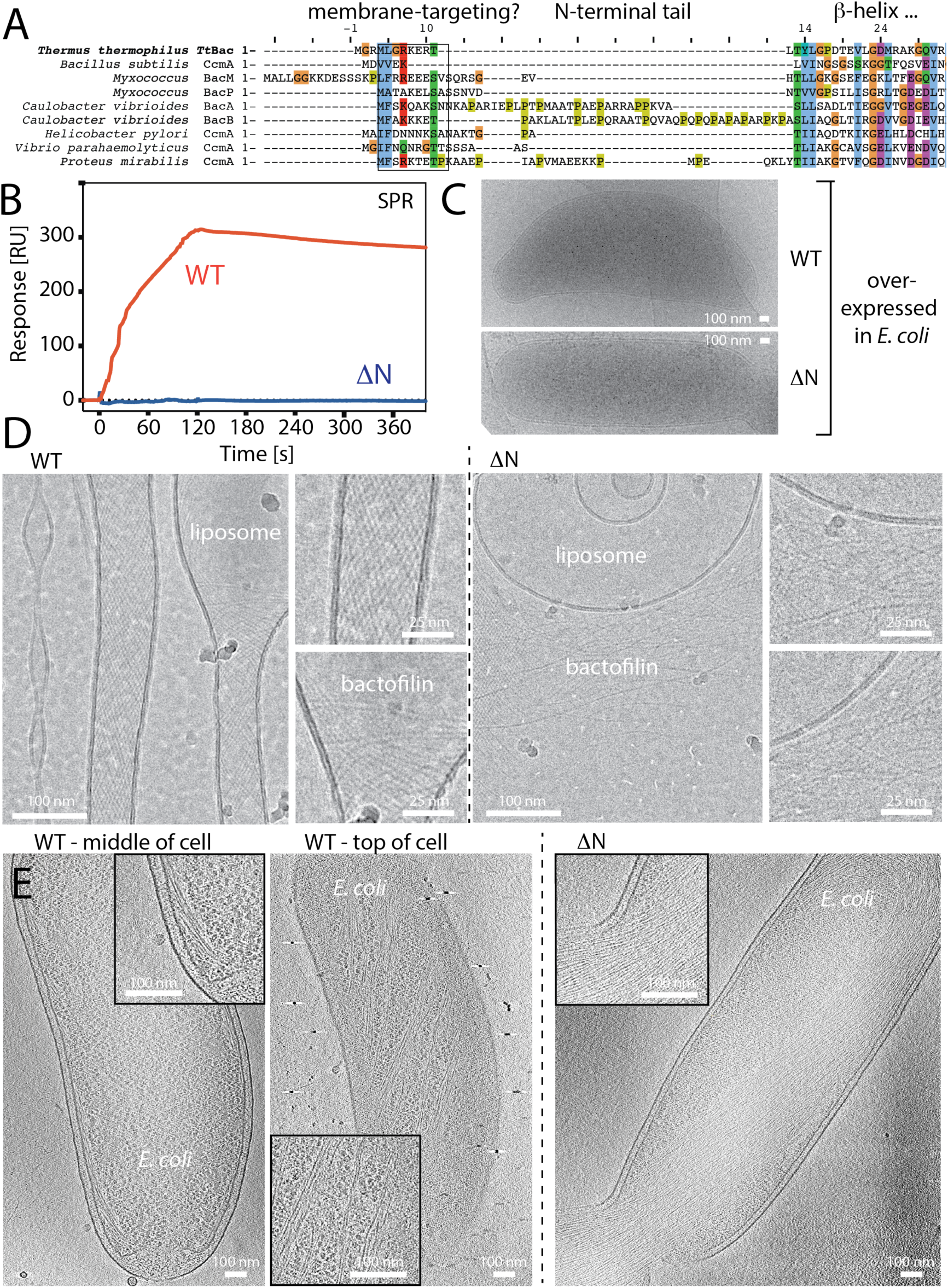
TtBac bactofilin binds to lipid membranes through its N-terminal tail. A) Multiple sequence alignment showing the N-terminal tails of bactofilins, including TtBac. A doublet of hydrophobic residues often followed by a positively charged residue is conserved across species. The fact that the N-terminal tails interact to form double helical filaments in our study (Figure 4E) and the occurance of unusual hydrophobicity in a disordered tail region prompted us to investigate membrane binding of TtBac *in vitro* and *in vivo*. B) Surface Plasmon Resonance (SPR) traces of TtBac-WT and ΔN-TtBac proteins when added to liposome-coated L1 sensor chips (GE Healthcare Life Sciences). The unmodified protein clearly shows binding, whereas the mutant protein lacking the N-terminal tails shows no signal. C) When overexpressed in *E. coli*, TtBac leads to a morphological phenotype, it causes the cells to become crescent shaped and this effect disappears when over-expressing ΔN-TtBac under the same conditions. D) When adding TtBac protein to pre-formed liposomes made from *E. coli* lipids, the protein was shown by subsequent cryo-EM imaging to deform and tubulate the liposomes. Note that single protofilaments appear to deform the liposomes from the outside (insets). When ΔN-TtBac was added to the same liposomes, no binding was observed and the liposomes were not deformed, consistent with the idea that the N-terminal tails of TtBac bactofilin, and presumably most bactofilins, are the membrane-targeting domains of these proteins. E) Investigating the overexpression system depicted in Figure 6C by electron cryotomography (cryo-ET). TtBac-WT (left two images) clearly formed long bundles going around the cell close to the membrane, indicative of direct binding to the inner membrane of *E. coli* as no putative accessory protein were expected to be present. When doing the same experiment with ΔN-TtBac, a very large bundle of filaments was observed going through the centre of the cell, not close to the membrane, and impeding cell division. These results again indicated that it is the N-terminal tails of bactofilins that facilitate direct membrane binding.

First we needed to show that TtBac does indeed bind to membranes directly. Because the protein is always filamentous, standard pelleting or flotation assays with liposomes proved difficult, so we used surface plasmon resonance (SPR) to investigate (Figure 6B). TtBac-WT showed a strong signal, indicating liposome binding. Equally, when overexpressing TtBac in *E. coli* cells, we noted a strong phenotype of bent cells, in our experience a hallmark of filament formation directly on the inner membrane on one side of the cell’s body (Figure 6C).

To demonstrate that the N-terminus of TtBac is necessary for membrane binding we produced an N-terminal deletion mutant ΔN-TtBac containing residues 11-123 and no other modifications (Supplementary Table T2). Both overexpression of the mutant in *E. coli* and incubation of the protein with liposomes and SPR showed that membrane binding and morphological changes were completely abolished (Figure 6B & C).

The binding of TtBac to membranes appears to include a demand for curvature, as seen by the morphological phenotype caused by overexpression of the full-length filamentous protein in *E. coli* (Figure 6C). Similarly, its interaction with liposomes *in vitro* is characterised by strong deformations or even tubulation (Figure 6D) and, again, this effect (and binding) is completely abolished when using ΔN-TtBac.

Finally, we also visualised TtBac-WT filaments in *E. coli* cells by cryo-ET, where they were visible as bundles circling the cell at an angle on or close to the inner membrane. The helicity (angle) confirms TtBac’s preference for curvature. When doing the same experiment with ΔN-TtBac, all membrane proximity was abolished and the filaments formed one large bundle in the cytoplasm (that was so large that it also inhibited cell division) (Figure 6E, left and right).

## Discussion

We found bactofilins in eukaryotes, although, as far as we could tell, restricted to Stramenopiles and Ascomycete fungi. Although more analysis will be needed, we believe that this is the result of horizontal gene transfer from prokaryotes into these organisms (McCarthy and Fitzpatrick, 2016; Richards et al., 2006). It will be interesting to determine the function of bactofilins in eukaryotes as well as in archaea where they appear to be more widespread.

Our structural work showed bactofilin TtBac from *Thermus thermophilus* to be closely related to BacA from *C. crescentus* (Shi et al., 2015); it polymerised into completely beta-helical protofilaments that have head-to-head (N-N) and tail-to-tail (C-C) interfaces. This arrangement produces non-polar protofilaments and because the two-fold axes at the N-N and C-C interface are nearly aligned, the filament twists only very slightly, ~5° per dimer of ~6 nm length. The unanticipated lack of polarity (Vasa et al., 2015; Zuckerman et al., 2015) makes the two ends of the filament equal and hence excludes all cytomotive mechanisms (Wagstaff and Löwe, 2018). Bactofilins should therefore be classed as cytoskeletal, along with MreB, DivIVA, SepF and MinCD, all of which form non-polar filaments in bacteria.

It is striking that the above list of cytoskeletal filaments contains exclusively cooperative filaments (Ghosal and Löwe, 2015) that bind directly to cell membranes from the inside. We showed here that bactofilins are no exception as they contain a conserved membrane targeting sequence (MTS) within their N-terminal tail.

From Figure 6D it is clear that the TtBac filaments form mostly single protofilaments when bound to membrane/liposomes, indicating and confirming that the N-terminal tails function to bind membrane and not each other, as in the double-helical filaments in our cryo-EM structure. We propose that the *bona fide* bactofilin filament is an almost straight, non-helical single protofilament, and that it is constitutively membrane bound (Supplementary Figure S6).

Both the tomography and microscopy (Figure 6C, D & E) demonstrated that TtBac preferred curvature, another feature that it shares with MreB (Hussain et al., 2018), DivIVA (Lenarcic et al., 2009) and SepF (Gündoğdu et al., 2011). Curvature preference in all the above filament systems seems to stem from the fact that they are not completely straight and this could also be a general mechanism of filament length restriction on flat, non-deformable membranes and/or curvature induction/sensing.

We firmly believe that bactofilins deserve greater attention as components of prokaryotic cytoskeletons. Armed with the analysis presented here demonstrating the widespread occurrence, biochemical stability, direct membrane binding, and non-polar filament architecture of these extraordinary proteins we hope that investigations will intensify into the roles of bactofilins in the cell biology of the organisms in which they are found. We anticipate further roles in morphogenesis and in processes where very stable membrane attachment is needed that cannot overlap with any of the other known systems. It also remains unclear how polymerisation, and with it membrane attachment, is regulated in cells and what role, if any, the C-terminal tails have.

## Materials & Methods

### Identification of bactofilins outside bacteria

We became interested in the possibility that bactofilins are present outside of the kingdom of bacteria after noting that PFAM (http://pfam.xfam.org) annotates several members of the PFAM 04519 (DUF 853, bactofilin) family in eukaryotes and archaea. This was investigated further by running a hmmsearch (HMMER3) (Eddy, 2011) with PFAM 04519 against the eukaryotic UniprotKB. The results included two taxonomic clusters of hits, one consisting of proteins found amongst the Stramenopiles, and the other of proteins within Ascomycete fungi. There were also hits in other eukaryotes, but these were either non-bactofilin repetitive sequences, or isolated hits within well-sequenced clades (in all these cases the best BLASTP matches were bacterial sequences). Clusters of hits within Stramenopiles and Ascomycetes, were aligned, HMMs were built and additional examples within UniprotKB were searched for. In both cases further examples were recovered from the relevant clades, but not outside them. To better understand the relationship between these putative eukaryotic bactofilins and the bacterial sequences, an alignment (using HMMALIGN) was made of putative bactofilin domains It comprised of bactofilins that are representative of sequence diversity within bacteria and archaea (found using the proGenomes set of representative prokaryotic genomes (Mende et al., 2017) and HMMSEARCH with PFAM 04519), and all of the putative eukaryotic bactofilins, including the taxonomic singletons, and the additional sequences from the putative clusters. From the alignment a phylogeny was constructed using using the FastTree algorithm, after selecting informative columns using GBlocks (Talavera and Castresana, 2007) (Supplementary Data D1). The phylogeny did not imply the existence of a common ancestor of eukaryotic bactofilins within eukaryotes. After inspecting alignments and trees three putative eukaryotic bactofilins were chosen to clone and express: one fungal protein (Q7RX79 from *Neurospora crassa*), and two Stramenopile proteins (D0N980 from *Phytophthora infestans*, and D7G0L1 from *Ectocarpus siliculosis*). Only D0N980 (PITG_07992) was successfully expressed and confirmed to be a bactofilin. It remains to be seen how many of the putative bactofilins outside the Oomycetes are polymerising bactofilins.

### Protein cloning, expression and purification

Amino acid sequences of all proteins used in this study are listed in Supplementary Table T2.

#### PiBac Oomycetes bactofilin

To obtain a plasmid that codes for full length bactofilin protein from *Phytophthora infestans* (PiBac), the gene PITG_07992 (UniProtKB, D0N980) was synthesised (GenScript), amplified and cloned into the pHis17 plasmid using Gibson Assembly (New England Biolabs), with a stop codon before the C-terminal hexahistidine tag on the plasmid. C41(DE3) *E. coli* cells (Lucigen) were transformed with the resulting plasmid by electroporation. 40 ml 2xTY media supplemented with 100 μg/ml ampicillin were inoculated with a single colony from the plate, and were grown at 200 rpm, 37 °C overnight. The pre-culture was then used to inoculate 4 L 2xTY media with 100 μg/ml ampicillin. After reaching an OD_600_ of 0.6-1.0 at 200 rpm, 37 °C, the expression was induced with 1 mM isopropyl β-D-thiogalactoside for 4 h at the same temperature, and the cells were harvested by centrifugation. For purification, the entire pellet was resuspended in 150 ml Buffer D (50 mM Tris, 200 NaCl, 1 mM TCEP, pH 8) supplemented with DNase I, RNase A (Sigma) and EDTA-free protease inhibitor tablets (Roche). Cells were lysed by sonication, and the lysate was cleared by centrifugation at 10,000 rpm in a 45 Ti rotor (Beckman) for 30 minutes at 4 °C, followed by another centrifugation at 20,000 rpm in a 45 Ti rotor for 30 minutes at 4 °C. The supernatant was supplemented with 2% v/w PEG 8000 prior to centrifugation again at 40,000 rpm in the 45 Ti rotor for 30 minutes at 4 °C. Each pellet was resuspended in 5 ml Buffer D supplemented with 0.3 g/ml caesium chloride and 1 % v/v Triton X-100, and the pellet was solubilised overnight at 4 °C. The solubilised mixture was centrifuged in a TLA 100.3 rotor (Beckman) at 50,000 rpm for 15 minutes at 4 °C, the pellet was discarded and the supernatant was centrifuged in a TLA 100.3 rotor at 80,000 rpm for 5 hours at 4 °C. The white layer near the bottom of the centrifuge tube was extracted, resuspended in 3 ml Buffer D supplemented with 0.3 g/ml caesium chloride before centrifugation again in a TLA 100.3 rotor at 80,000 rpm overnight at 4 °C. The white layer containing filaments near the bottom of the centrifuge tube was extracted again and examined using negative stain electron microscopy (see below). Identity of the protein was confirmed by SDS-PAGE.

#### H_6_-TtBac 1-123

To obtain an *E. coli* expression plasmid for N-terminally His_6_-tagged full-length bactofilin protein from *Thermus thermophilus* (TtBac, MGSSHHHHHH-1-123), the gene coding for protein WP_011173792.1 (NCBI) was amplified from genomic DNA and was cloned into vector pET15b using Gibson Assembly (New England Biolabs). C41(DE3) *E. coli* cells (Lucigen) were transformed by electroporation. 60 ml 2xTY media supplemented with 100 μg/ml ampicillin were inoculated with a single colony from the plate, and were grown at 200 rpm, 37 °C overnight. The culture was then used to inoculate 6 L 2xTY media with 100 μg/ml ampicillin. After reaching an OD_600_ of 0.6-1.0 at 200 rpm, 37 °C, protein expression was induced with 1 mM isopropyl β-D-thiogalactoside for 4 h at the same temperature, and the cells were harvested by centrifugation. For purification, the entire pellet was resuspended in 200 ml Buffer A (50 mM Tris, 6 M guanidinium chloride, 1 mM TCEP, pH 7). Cells were disrupted at 25 kPSI in a cell disruptor (Constant Systems), and the lysate was cleared by centrifugation at 40,000 rpm in a 45 Ti rotor (Beckman) for 30 minutes at 25 °C. The cleared lysate was loaded onto a 5 ml HisTrap HP column (GE Healthcare), and was washed with stepwise increases of imidazole in Buffer A: 0, 20, 50, 200, 500 and 1000 mM. Eluted fractions was analysed by SDS-PAGE and those containing TtBac (mostly at 50 mM imidazole) were pooled and concentrated in Centriprep concentrators (10 kDa MWCO, Millipore) to 10 mg/ml. The H**_6_**-TtBac protein was refolded by a single step dialysis process into Buffer B (50 mM CAPS, 200 NaCl, 1 mM TCEP, pH 11) overnight at room temperature. The protein was stored at 4 °C.

#### TtBac-WT 1-123

To obtain an *E. coli* expression plasmid for untagged, full-length bactofilin protein from *Thermus thermophilus* (TtBac, 1-123), the gene coding for protein WP_011173792.1 (NCBI) was amplified from the above H**_6_**-TtBac pET15b plasmid and was cloned into plasmid pHis17 using Gibson Assembly (New England Biolabs) with a stop codon before the C-terminal tag on the plasmid. The plasmid encoding untagged TtBac was used to transform C41(DE3) *E. coli* cells (Lucigen) by electroporation. 40 ml 2xTY media supplemented with 100 μg/ml ampicillin were inoculated with a single colony from the plate, and were grown at 200 rpm, 37 °C overnight. The culture was then used to inoculate 4 L 2xTY media with 100 μg/ml ampicillin. After reaching an OD_600_ of 0.6-1.0 at 200 rpm, 37 °C, the expression was induced with 1 mM isopropyl β-D-thiogalactoside for 4 h at the same temperature, and the cells were harvested by centrifugation. Purification was achieved by modifying a previous protocol (Koch et al., 2011). For purification, the entire pellet was resuspended in 150 Buffer B (50 mM CAPS, 200 NaCl, 1 mM TCEP, pH 11) supplemented with DNase I, RNase A (Sigma) and EDTA-free protease inhibitor tablets (Roche). Cells were lysed by sonication, and the lysate was cleared by centrifugation at 10,000 rpm in a 45 Ti rotor (Beckman) for 30 minutes at 4 °C, followed by another centrifugation at 20,000 rpm in the 45 Ti rotor for 30 minutes at 4 °C. The supernatant was supplemented with 2% v/w PEG 8000 prior to centrifugation at 40,000 rpm in the 45 Ti rotor for 30 minutes at 4 °C. Each pellet was resuspended in 5 ml Buffer B supplemented with 0.3 g/ml caesium chloride and 1 % v/v Triton X-100, and the pellet was solubilised overnight at 4 °C. The mixture was centrifuged in a TLA 100.3 rotor (Beckman) at 50,000 rpm for 15 minutes at 4 °C, the pellet was discarded and the supernatant was centrifuged in the TLA 100.3 rotor at 80,000 rpm for 5 hours at 4 °C. The white layer containing TtBac filaments near the bottom of the tube were extracted, resuspended in 3 ml Buffer B supplemented with 0.3 g/ml caesium chloride before centrifugation in the TLA 100.3 rotor at 80,000 rpm overnight at 4 °C. The layer containing TtBac filaments near the bottom of the centrifuge tube was extracted, and diluted with an equal volume of Buffer C (50 mM CAPS, 100 mM NaCl, 1 mM TCEP, 1 mM EDTA, pH 11) before being loaded onto a Superose 6 10/300 size exclusion column (GE Healthcare), equilibrated in Buffer C. The fractions containing TtBac filaments (in the void volume of the column) were pooled, and centrifuged in a TLA 100.3 rotor (Beckman) at 80,000 rpm for 30 minutes at 4 °C. The TtBac filaments formed a transparent pellet at the bottom of the tube, and were resuspended in Buffer C to 1-2 mg/ml and stored at 4 °C.

#### Nanobody NB4-mut2

Purified and refolded H**_6_**-TtBac filaments were sent to VIB’s Nanobody Core for commercial camelid nanobody generation and selection. Frozen bacterial cultures containing individual nanobody genes in the pMECS vector were delivered by VIB. Genes encoding for each nanobody were amplified using colony PCR and cloned into the pHEN6c expression vector (VIB) using Gibson Assembly (New England Biolabs). The resulting constructs using pHEN6c encoded for PelB(leader)-nanobody-SSHHHHHH proteins.

For expression, the pHEN6c plasmids encoding for the nanobody proteins were used to transform WK6 *E. coli* cells (VIB, essential) by electroporation. 20 ml 2xTY media supplemented with 100 μg/ml ampicillin were inoculated with a single colony from the plate, and were grown at 200 rpm, 37 °C overnight. The culture was then used to inoculate 2 L 2xTY media with 100 μg/ml ampicillin. After reaching an OD_600_ of 0.6-1.0 at 200 rpm, 37 °C, the expression was induced with 1 mM isopropyl β-D-thiogalactoside for 4 h at the same temperature, and the cells were harvested by centrifugation.

The purification protocol followed recommendations from VIB. Since the plasmids encoded the PelB leader sequence, nanobody proteins were secreted into the periplasm. For purification, the entire pellet from 2 L culture was resuspended in 24 ml TES buffer (200 mM Tris, 500 mM sucrose, 0.5 mM EDTA, pH 8), with shaking at 4 °C for 1 hour. The mixture was supplemented with 36 ml TES/4 buffer (TES buffer diluted 4 times in water) and was mixed at 4 °C for 1 hour. The suspension was centrifuged in a 45 Ti rotor (Beckman) for 30 minutes at 40,000 rpm at 4 °C. The supernatant containing the periplasmic extract was sonicated before being loaded onto a 5 ml HisTrap HP column (GE Healthcare). The column was washed with stepwise increases of imidazole in buffer E (50 mM Tris, 200 mM NaCl, 1 mM TCEP, pH 8): 3, 200, 500 and 1000 mM. Elutions were analysed by SDS-PAGE and the fractions containing nanobodies (mostly at 200 mM imidazole) were pooled and concentrated in Centriprep concentrators (10 kDa MWCO, Millipore) to around 6 mg/ml. The concentrated proteins were buffer exchanged using a PD-10 desalting column (GE Healthcare) equilibrated with Buffer C (50 mM CAPS, 100 mM NaCl, 1 mM TCEP, 1 mM EDTA, pH 11) using the spin protocol.

To solve the bundling issue when nanobody proteins were added to bactofilin filaments, mutants were made to try to change selected surface amino acid residues into aspartic acids using nanobody 4 (NB4) as template, that had been selected by negative stain EM of TtBac and nanobody equimolar mixtures. The gene encoding NB4 mutant 2 (L13S, Q15D, K45D, K66D) was synthesised (GenScript), amplified and cloned into the pHEN6c expression vector. Nanobody mutant proteins were expressed and purified using the same protocol as the non-mutated nanobody proteins. The final concentration of NB4 mutant 2 was 6.8 mg/ml.

#### ΔN-TtBac(F105R)-H_6_

The required coding region was amplified from the H_6_-TtBac construct by PCR and was cloned into the plasmid pHis17 using Gibson Assembly (New England Biolabs), resulting in a C-termin tag: GSHHHHHH. The point mutation was introduced by Q5 mutagenesis (NEB). The resulting plasmid was used to transform C41(DE3) *E. coli* cells (Lucigen) by electroporation. 12L cultures in 2xTY were inoculated and induced with 1 mM IPTG at an OD600 of 0.6 and further grown for 6 h at 37 °C. After harvesting of the cells, the pellet was dissolved in Buffer F (50 mM Tris/HCl, 200 mM NaCl, pH 7.5) and lysed by cell disruption at 35 kPSI (Constant Systems). The lysate was cleared by ultracentrifugation (2 h at 35,000 rpm in a 45 Ti rotor) and loaded onto a 5 ml HisTrap column. The bound fraction was eluted with 0.6 M imidazole, pH 7.0 and further purified by size exclusion on a Sephacryl S300 16/60 column (GE Healthcare) in Buffer G (20 mM CHES/NaOH, 250 mM NaCl, pH 9.5). Fractions were analysed by SDS-PAGE and the fractions containing ΔN-TtBac(F105R)-H_6_ were pooled and concentrated in Centriprep concentrators (10 kDa MWCO, Millipore) to around 12 mg/ml.

#### ΔN-TtBac 11-123

The required coding region was amplified from the H_6_-TtBac construct by PCR and was cloned into the plasmid pHis17 using Gibson Assembly (New England Biolabs), The plasmid was used to transform C41(DE3) *E. coli* cells (Lucigen) by electroporation. 12 L cultures in 2xTY were induced with 1 mM IPTG at an OD600 of 0.6 and further grown for 4 h at 37 °C. After harvesting of the cells, the cells were dissolved in the same buffer as for TtBac-WT 1-123 and spun for 30 min at 9k rpm in a Ti45 rotor (Beckman). The pellet was dissolved in the same buffer and from here onwards the same protocol as for TtBac-WT 1-123 was used.

### Electron microscopy: negative stain

Continuous carbon grids were purchased from EMS (Electron Microscopy Sciences). After glow discharging, 3 µl of sample were applied, blotted and stained with fresh 2% uranyl acetate solution. Uranyl acetate was applied 1-3 times with wait times of up to 60 seconds. After air-drying, grids were imaged in an FEI F20 electron microscope equipped with a Falcon 2 detector, operated at room temperature.

### Electron microscopy: cryo-EM, helical reconstruction of TtBac-WT and with nanobody NB4-mut2

Initially, refolded and His_6_-tagged TtBac protein (H**_6_**-TtBac) was investigated using electron cryomicroscopy (cryo-EM) with the aim to obtain an atomic model of the filament. For this, 2 μl 60 μM H**_6_**-TtBac in Buffer C were added onto freshly glow-discharged Quantifoil Cu/Rh R2/2 holey carbon 200 mesh grids (Quantifoil). The grids were blotted for 2.5 s with a blotting force of −15, a drain time of 0.5 s, and were flash frozen in liquid nitrogen-cooled liquid ethane using an FEI Vitrobot Mark IV. The Vitrobot chamber was set to 10 °C and 100 % humidity.

Grids were imaged at 300 kV using a FEI Tecnai Polara G2 or Titan Krios microscope using a Falcon III detector (one dataset was collected at eBIC, Harwell, UK), using pixel sizes between 1.0 to 1.4 Å and average total doses of about 40 e^−^/Å^2^, distributed over 40-70 frames. Images were motion corrected using MOTIONCOR2 (Zheng et al., 2017) and CTF corrected using GCTF (Zhang, 2016). 2D classification analysis in RELION (Scheres, 2012) revealed that the filaments had varying widths, most likely caused by varying protofilament numbers that ranged from 2-4. We then switched to untagged and natively-purified TtBac-WT material, using the same vitrification and imaging conditions as above. The data showed very similar variations in protofilament number but it was possible to discern the 4.7 Å axial repeat caused by the beta-stacking of the subunits in 2D classes obtained by processing in RELION, however, it proved impossible to go beyond 5 Å resolution when performing reconstructions in RELION, most likely because the filaments are smooth and the starts and ends of each subunit could not be determined during RELION’s 3D refinement procedure. In order to overcome this issue, a nanobody was obtained, NB4, that showed very clear binding to the filaments as it changed their appearance and also led to a strong reduction in filaments with more than 2 protofilaments. Unfortunately, the filaments coated with NB4 tended to bundle heavily, impeding further analysis. Hence four mutations were introduced, L13S, Q15D, K45D and K66S, yielding NB4-mut2 nanobody that significantly reduced bundling and enabled image analysis and helical reconstruction of TtBac-WT bactofilin to near-atomic resolution. For the cryo-EM grid preparation with the nanobody, purified nanobody protein was added to TtBac filaments at a ratio of 1.2 : 1 (nanobody:filament). The final dataset used is summarised in Supplementary Table T1. It was collected on a FEI Titan Krios microscope at 300 kV, using a Falcon III detector in electron counting mode with an estimated pixel size of 1.07 Å/pixel. 2130 good images were selected after MOTIONCOR2 and GTCF and from those around 456,000 helical segments were picked in RELION, 57 Å apart along the filament axis. 2D classification in RELION selected 346,000 good helical segments and these were used for helical reconstruction in RELION with helical parameters twist = 4.73° and rise = 57.48 Å (He and Scheres, 2017). Particle polishing and post-processing followed standard RELION procedures and led to a final map with a resolution determined from gold-standard two halves FSC calculations (FSC = 0.143) of 3.6 Å (Rosenthal and Henderson, 2003). When inspecting the map this value hides that fact that the map has very anisotropic resolution, most likely caused by the dominant 4.7 Å beta-stacking repeat along the filament axis and has poor resolution perpendicular to the filament axis. The previously determined *B. subtilis* BacA ssNMR structure (PDB 2N3D) (Shi et al., 2015) was homology modelled into TtBac using the SWISSMODEL server and placed in the cryo-EM map, as was a SWISSMODEL of the NB4-mut2 nanobody. Placing these models produced excellent fit but the anisotropy of the map made atomic refinement difficult.

After the crystal structure of ΔN-TtBac(F105R)-H_6_ had been solved during the course of this study, a much better atomic model became available. And after performing signal substraction of the nanobody density from all 346,000 helical segment images in RELION (Bai et al., 2015), an improved cryo-EM map of the two TtBac protofilaments was obtained using RELION with helical parameters twist = 4.89° and rise = 57.46 Å. Several cycles of manual model adjustment in MAIN and refinement with PHENIX.real_space_refine (with additional secondary structure restraints) (Afonine et al., 2018) produced the final model (Supplementary Table 1). Resolution was estimated to be 3.4 Å from half-map FSC analysis in RELION but FSC analysis against the atomic model yielded a lower value of 4.2 Å, caused by the strong resolution anisotropy of the map. The atomic coordinates have been deposited in the Protein Data Bank (PDB) with accession number 6RIB and the subtracted cryo-EM map has been deposited in EMDB with accession code EMD-4887.

### Evolutionary sequence coupling analysis of filament formation

A large set of bactofilin sequences was obtained by searching the UniprotKB with the PFAM HMM PF04519 using HMMSEARCH with defaults. These 20,746 sequences were aligned to PF04519 using HMMALIGN. Inserts relative to the HMM were removed, and sequences with more than 5 % gaps relative to the HMM were discarded using the dcaTools package (gitlab.com/ducciomalinverni/dcaTools/), leaving 12,646 sequences. This alignment was used to infer a Direct Coupling Analysis (DCA) model of the bactofilin family using lbsDCA (gitlab.com/ducciomalinverni/lbsDCA/), an implementation of the asymmetric Pseudo-Likelihood Method for DCA (Ekeberg et al., 2013). lbsDCA reported an effective number of sequences after re-weighting with a 90 % identity cutoff of 5,770.4. The average-product corrected Frobenius norm DCA scores reported were scaled arbitrarily for plotting after smoothing of the heatmap with a 3×3 Gaussian kernel.

### Crystal structure determination of ΔN-TtBac(F105R)-H_6_

ΔN-TtBac(F105R)-H_6_ protein was produced as described above and selenomethionine-substituted protein (SeMet) was obtained using published protocols (van den Ent et al., 1999) with the same subsequent purification protocol as for the native protein, with the sole change that all buffers contained reducing agent TCEP at 1 mM. Initial crystallisation conditions were obtained using LMB’s in-house high-throughput crystallisation facility (Stock et al., 2005). All crystals were produced in MRC 2-drop crystallisation plates using 100 + 100 nl sitting drop setups and crystallisation experiments were performed at 19 °C. SeMet ΔN-TtBac(F105R)-H_6_ was crystallised using reservoir solution containing 6.7-7.3 v/v % 2-propanol, 0.17-0.19 M lithium sulfate, 0.1 M phosphate citrate, pH 3.7 and 0.4 M ammonium acetate as an additive. The crystals were cryo-cooled using 30% glycerol in reservoir solution. The crystals belonged to spacegroup I2_1_2_1_2_1_ and showed a very large unit cell (Supplementary Table T1). Three 360° datasets were collected at beamline I03 (Diamond Light Source, Harwell, UK) and merged together using CCP4 programs (Winn et al., 2011) resulting in very high multiplicity and extending to about 4.0 Å resolution. 20-30 selenium sites were readily identified using SHELXD (Sheldrick, 2008) and subsequent phasing with PHASER (Read and McCoy, 2011) and NCS averaging (operators deduced from SeMet sites within PHENIX) using DM resulted in interpretable electron density maps. It was determined that the number of recognisable subunits was 32 and both phasing and NCS averaging were adjusted to take this into account to obtain a final electron density map. The preliminary atomic model obtained from cryo-EM maps at 4-5 Å resolution was fitted manually, guided by the position of the single SeMet25 residue and its anomalous signal, and it was recognised that residues A101-G112 had become disordered, presumably caused by the F105R mutation used in the construct. Analysis then switched to the native protein. Native ΔN-TtBac(F105R)-H_6_ was crystallised using reservoir solutions as above and the crystals were cryo-cooled as above. The crystals again belonged to spacegroup I2_1_2_1_2_1_ with only slightly different cell constants. The crystals were isomorphous enough to enable simple rigid body refinement of all 32 chains since molecular replacement was not possible, presumably because of the smooth appearance of the protofilaments that makes it difficult to discerns the starts and ends of each subunit, very similar to the initial problems with cryo-EM image reconstruction. The structure was rebuilt manually in MAIN (Turk, 2013) and refined with Phenix.refine (Adams et al., 2010) (using NCS restraints and also real space refinement) for several cycles. Final statistics are summarised in Supplementary Table T1 and the resulting coordinates haven been deposited in the Protein Data Bank (PDB) with accession code 6RIA.

### Surface Plasmon Resonance

Surface plasmon resonance (SPR) data were collected using a BIAcore T200 instrument using a L1 Sensor Chip (GE Healthcare). Both reference and ligand channels were equilibrated in 50 mM CAPS, pH 8.0, 100 mM NaCl, 4 mM TCEP at 25 °C. Liposomes were prepared with *E. coli* Total Lipid Extract (Avanti Polar Lipids) using freeze-thaw cycles followed by sonication in buffer. Liposomes at a lipid concentration of 3 mg/ml were captured onto the ligand surface at 10 µL/min to a level of ~ 1200 RU. To prevent non-specific binding the surfaces were passivated by injections of 0.2 mg/ml BSA with 1 mg/ml NSB (GE Healthcare Life Sciences) for 120 s prior to a 120 s injection at 30 µl/min of either TtBac-WT, ΔN-TtBac at 10 µM or buffer followed by dissociation for 300 s. After each measurement the surfaces were regenerated with a 30 s injection of 20 mM CHAPS. Data were doubly-referenced by subtraction of the reference channel data and from injections of buffer alone.

### Bactofilin binding to liposomes

Liposomes were prepared with *E. coli* total lipid extract (Avanti Polar Lipids) using freeze-thaw cycles followed by sonication in buffer D (50 mM CAPS, 100 mM NaCl, 4 mM TCEP, pH 8.0). Images of liposomes together with native TtBac-WT and ΔN-TtBac were obtained by mixing 120 μM protein in buffer C with 2 mg/ml liposomes in buffer D. Of this mixture, 3 μl were applied onto freshly glow-discharged Quantifoil Cu/Rh R2/2 holey carbon 200 mesh grids (Quantifoil). The grids were blotted for 4 seconds with a blotting force of −15, a drain time of 0.5 s, and were flash frozen in liquid-nitrogen-cooled liquid ethane using an Vitrobot Mark IV (FEI). The Vitrobot chamber was set to 10 °C and 100 % humidity. Grids were imaged in an FEI F20 electron microscope equipped with a Falcon 2 detector, operated at cryogenic temperature.

### Bactofilin overexpression in E. coli and electron tomography

Cells expressing TtBac-WT or ΔN-TtBac were mixed with 10 nm protein-A gold fiducials and plunge frozen on Quantifoil R2/2 holey carbon grids using a Vitrobot Mark IV (FEI). Tomography data were collected on a Titan Krios electron microscope equipped with a Quantum imaging filter and K2 direct detector (both Gatan). Tilt series were acquired using SerialEM (Mastronarde, 2005) from 0° to ± 60° using a grouped dose symmetric tilt scheme (Hagen et al., 2017) with a 2° increment and a total dose of 160 e^−^/Å^2^. The pixel size was 5.44 Å, the target defocus 8 μm and the slit width of the energy filter 20 eV. Tomograms were reconstructed in IMOD (Kremer et al., 1996), using the SIRT algorithm.

## Acknowledgements

We would like to thank Howard S. Judelson (UC Riverside) for discussions regarding Oomycetes. We acknowledge Diamond Light Source for access and support of the cryo-EM facilities at the UK’s national Electron Bio-imaging Centre (eBIC), funded by the Wellcome Trust, MRC and BBRSC. We would like to thank Minmin Yu (MRC-LMB) for help with synchrotron data collection and staff at beamline I03 (Diamond Light Source, Harvell, UK) for excellent service provision. This work was funded by the Medical Research Council (U105184326 to JL) and the Wellcome Trust (202754/Z/16/Z to JL). James Wagstaff and Xian Deng were also supported by studentships from the Boehringer Ingelheim Fonds.

## References

Adams, P. D., Afonine, P. V., Bunkóczi, G., Chen, V. B., Davis, I. W., Echols, N., Headd, J. J., Hung, L. W., Kapral, G. J., Grosse-Kunstleve, R. W., McCoy, A. J., Moriarty, N. W., Oeffner, R., Read, R. J., Richardson, D. C., Richardson, J. S., Terwilliger, T. C., and Zwart, P. H. (2010). PHENIX: a comprehensive Python-based system for macromolecular structure solution. Acta Crystallogr D Biol Crystallogr 66, 213–221.

Afonine, P. V., Poon, B. K., Read, R. J., Sobolev, O. V., Terwilliger, T. C., Urzhumtsev, A., and Adams, P. D. (2018). Real-space refinement in PHENIX for cryo-EM and crystallography. Acta Crystallogr D Struct Biol 74, 531–544.

Ah-Fong, A. M., Kim, K. S., and Judelson, H. S. (2017). RNA-seq of life stages of the oomycete Phytophthora infestans reveals dynamic changes in metabolic, signal transduction, and pathogenesis genes and a major role for calcium signaling in development. BMC Genomics 18, 198.

Amos, L. A., and Löwe, J. (2017). Overview of the Diverse Roles of Bacterial and Archaeal Cytoskeletons. Subcell Biochem 84, 1–26.

Bai, X. C., Rajendra, E., Yang, G., Shi, Y., and Scheres, S. H. (2015). Sampling the conformational space of the catalytic subunit of human γ-secretase. Elife 4,

Blair, K. M., Mears, K. S., Taylor, J. A., Fero, J., Jones, L. A., Gafken, P. R., Whitney, J. C., and Salama, N. R. (2018). The Helicobacter pylori cell shape promoting protein Csd5 interacts with the cell wall, MurF, and the bacterial cytoskeleton. Mol Microbiol 110, 114–127.

Bulyha, I., Lindow, S., Lin, L., Bolte, K., Wuichet, K., Kahnt, J., van der Does, C., Thanbichler, M., and Søgaard-Andersen, L. (2013). Two small GTPases act in concert with the bactofilin cytoskeleton to regulate dynamic bacterial cell polarity. Dev Cell 25, 119–131.

Derelle, R., López-García, P., Timpano, H., and Moreira, D. (2016). A Phylogenomic Framework to Study the Diversity and Evolution of Stramenopiles (=Heterokonts). Mol Biol Evol 33, 2890–2898.

Eddy, S. R. (2011). Accelerated Profile HMM Searches. PLoS Comput Biol 7, e1002195.

Ekeberg, M., Lövkvist, C., Lan, Y., Weigt, M., and Aurell, E. (2013). Improved contact prediction in proteins: using pseudolikelihoods to infer Potts models. Phys Rev E Stat Nonlin Soft Matter Phys 87, 0012707.

El Andari, J., Altegoer, F., Bange, G., and Graumann, P. L. (2015). Bacillus subtilis Bactofilins Are Essential for Flagellar Hook- and Filament Assembly and Dynamically Localize into Structures of Less than 100 nm Diameter underneath the Cell Membrane. PLoS One 10, e0141546.

El-Gebali, S., Mistry, J., Bateman, A., Eddy, S. R., Luciani, A., Potter, S. C., Qureshi, M., Richardson, L. J., Salazar, G. A., Smart, A., Sonnhammer, E. L. L., Hirsh, L., Paladin, L., Piovesan, D., Tosatto, S. C. E., and Finn, R. D. (2019). The Pfam protein families database in 2019. Nucleic Acids Res 47, D427–D432.

Ghosal, D., and Löwe, J. (2015). Collaborative protein filaments. EMBO J 34, 2312–2320.

Gode-Potratz, C. J., Kustusch, R. J., Breheny, P. J., Weiss, D. S., and McCarter, L. L. (2011). Surface sensing in Vibrio parahaemolyticus triggers a programme of gene expression that promotes colonization and virulence. Mol Microbiol 79, 240–263.

Gündoğdu, M. E., Kawai, Y., Pavlendova, N., Ogasawara, N., Errington, J., Scheffers, D. J., and Hamoen, L. W. (2011). Large ring polymers align FtsZ polymers for normal septum formation. EMBO J 30, 617–626.

Hagen, W. J. H., Wan, W., and Briggs, J. A. G. (2017). Implementation of a cryo-electron tomography tilt-scheme optimized for high resolution subtomogram averaging. J Struct Biol 197, 191–198.

Hay, N. A., Tipper, D. J., Gygi, D., and Hughes, C. (1999). A novel membrane protein influencing cell shape and multicellular swarming of Proteus mirabilis. J Bacteriol 181, 2008–2016.

He, S., and Scheres, S. H. W. (2017). Helical reconstruction in RELION. J Struct Biol 198, 163–176.

Hussain, S., Wivagg, C. N., Szwedziak, P., Wong, F., Schaefer, K., Izoré, T., Renner, L. D., Holmes, M. J., Sun, Y., Bisson-Filho, A. W., Walker, S., Amir, A., Löwe, J., and Garner, E. C. (2018). MreB filaments align along greatest principal membrane curvature to orient cell wall synthesis. Elife 7,

Jackson, K. M., Schwartz, C., Wachter, J., Rosa, P. A., and Stewart, P. E. (2018). A widely conserved bacterial cytoskeletal component influences unique helical shape and motility of the spirochete Leptospira biflexa. Mol Microbiol 108, 77–89.

Kassem, M. M., Wang, Y., Boomsma, W., and Lindorff-Larsen, K. (2016). Structure of the Bacterial Cytoskeleton Protein Bactofilin by NMR Chemical Shifts and Sequence Variation. Biophys J 110, 2342–2348.

Koch, M. K., McHugh, C. A., and Hoiczyk, E. (2011). BacM, an N-terminally processed bactofilin of Myxococcus xanthus, is crucial for proper cell shape. Mol Microbiol 80, 1031–1051.

Kremer, J. R., Mastronarde, D. N., and McIntosh, J. R. (1996). Computer visualization of three-dimensional image data using IMOD. J Struct Biol 116, 71–76.

Kühn, J., Briegel, A., Mörschel, E., Kahnt, J., Leser, K., Wick, S., Jensen, G. J., and Thanbichler, M. (2010). Bactofilins, a ubiquitous class of cytoskeletal proteins mediating polar localization of a cell wall synthase in Caulobacter crescentus. EMBO J 29, 327–339.

Lenarcic, R., Halbedel, S., Visser, L., Shaw, M., Wu, L. J., Errington, J., Marenduzzo, D., and Hamoen, L. W. (2009). Localisation of DivIVA by targeting to negatively curved membranes. EMBO J 28, 2272–2282.

Lin, L., Osorio Valeriano, M., Harms, A., Søgaard-Andersen, L., and Thanbichler, M. (2017). Bactofilin-mediated organization of the ParABS chromosome segregation system in Myxococcus xanthus. Nat Commun 8, 1817.

Lin, L., and Thanbichler, M. (2013). Nucleotide-independent cytoskeletal scaffolds in bacteria. Cytoskeleton (Hoboken) 70, 409–423.

Mastronarde, D. N. (2005). Automated electron microscope tomography using robust prediction of specimen movements. J Struct Biol 152, 36–51.

McCarthy, C. G., and Fitzpatrick, D. A. (2016). Systematic Search for Evidence of Interdomain Horizontal Gene Transfer from Prokaryotes to Oomycete Lineages. mSphere 1,

Mende, D. R., Letunic, I., Huerta-Cepas, J., Li, S. S., Forslund, K., Sunagawa, S., and Bork, P. (2017). proGenomes: a resource for consistent functional and taxonomic annotations of prokaryotic genomes. Nucleic Acids Res 45, D529–D534.

Mendler, K., Chen, H., Parks, D. H., Hug, L. A., and Doxey, A. AnnoTree: visualization and exploration of a functionally annotated microbial tree of life. BioRxiv 463455,

Michie, K. A., and Löwe, J. (2006). Dynamic filaments of the bacterial cytoskeleton. Annu Rev Biochem 75, 467–492.

Miraldi, E. R., Thomas, P. J., and Romberg, L. (2008). Allosteric models for cooperative polymerization of linear polymers. Biophys J 95, 2470–2486.

Parks, D. H., Chuvochina, M., Waite, D. W., Rinke, C., Skarshewski, A., Chaumeil, P. A., and Hugenholtz, P. (2018). A standardized bacterial taxonomy based on genome phylogeny substantially revises the tree of life. Nat Biotechnol 36, 996–1004.

Pichoff, S., and Lutkenhaus, J. (2005). Tethering the Z ring to the membrane through a conserved membrane targeting sequence in FtsA. Mol Microbiol 55, 1722–1734.

Rajagopala, S. V., Titz, B., Goll, J., Parrish, J. R., Wohlbold, K., McKevitt, M. T., Palzkill, T., Mori, H., Finley, R. L., and Uetz, P. (2007). The protein network of bacterial motility. Mol Syst Biol 3, 128.

Read, R. J., and McCoy, A. J. (2011). Using SAD data in Phaser. Acta Crystallogr D Biol Crystallogr 67, 338–344.

Richards, T. A., Dacks, J. B., Jenkinson, J. M., Thornton, C. R., and Talbot, N. J. (2006). Evolution of filamentous plant pathogens: gene exchange across eukaryotic kingdoms. Curr Biol 16, 1857–1864.

Rosenthal, P. B., and Henderson, R. (2003). Optimal determination of particle orientation, absolute hand, and contrast loss in single-particle electron cryomicroscopy. J Mol Biol 333, 721–745.

Salje, J., van den Ent, F., de Boer, P., and Löwe, J. (2011). Direct membrane binding by bacterial actin MreB. Mol Cell 43, 478–487.

Scheres, S. H. (2012). RELION: implementation of a Bayesian approach to cryo-EM structure determination. J Struct Biol 180, 519–530.

Sheldrick, G. M. (2008). A short history of SHELX. Acta Crystallogr A 64, 112–122.

Shi, C., Fricke, P., Lin, L., Chevelkov, V., Wegstroth, M., Giller, K., Becker, S., Thanbichler, M., and Lange, A. (2015). Atomic-resolution structure of cytoskeletal bactofilin by solid-state NMR. Sci Adv 1, e1501087.

Stock, D., Perisic, O., and Löwe, J. (2005). Robotic nanolitre protein crystallisation at the MRC Laboratory of Molecular Biology. Prog Biophys Mol Biol 88, 311–327.

Sycuro, L. K., Pincus, Z., Gutierrez, K. D., Biboy, J., Stern, C. A., Vollmer, W., and Salama, N. R. (2010). Peptidoglycan crosslinking relaxation promotes Helicobacter pylori’s helical shape and stomach colonization. Cell 141, 822–833.

Szeto, T. H., Rowland, S. L., Rothfield, L. I., and King, G. F. (2002). Membrane localization of MinD is mediated by a C-terminal motif that is conserved across eubacteria, archaea, and chloroplasts. Proc Natl Acad Sci U S A 99, 15693–15698.

Talavera, G., and Castresana, J. (2007). Improvement of phylogenies after removing divergent and ambiguously aligned blocks from protein sequence alignments. Syst Biol 56, 564–577.

Turk, D. (2013). MAIN software for density averaging, model building, structure refinement and validation. Acta Crystallogr D Biol Crystallogr 69, 1342–1357.

van den Ent, F., Lockhart, A., Kendrick-Jones, J., and Löwe, J. (1999). Crystal structure of the N-terminal domain of MukB: a protein involved in chromosome partitioning. Structure 7, 1181–1187.

Vasa, S., Lin, L., Shi, C., Habenstein, B., Riedel, D., Kühn, J., Thanbichler, M., and Lange, A. (2015). β-Helical architecture of cytoskeletal bactofilin filaments revealed by solid-state NMR. Proc Natl Acad Sci U S A 112, E127–36.

Wagstaff, J., and Löwe, J. (2018). Prokaryotic cytoskeletons: protein filaments organizing small cells. Nat Rev Microbiol 16, 187–201.

Wagstaff, J. M., Tsim, M., Oliva, M. A., García-Sanchez, A., Kureisaite-Ciziene, D., Andreu, J. M., and Löwe, J. (2017). A Polymerization-Associated Structural Switch in FtsZ That Enables Treadmilling of Model Filaments. MBio 8,

Winn, M. D., Ballard, C. C., Cowtan, K. D., Dodson, E. J., Emsley, P., Evans, P. R., Keegan, R. M., Krissinel, E. B., Leslie, A. G., McCoy, A., McNicholas, S. J., Murshudov, G. N., Pannu, N. S., Potterton, E. A., Powell, H. R., Read, R. J., Vagin, A., and Wilson, K. S. (2011). Overview of the CCP4 suite and current developments. Acta Crystallogr D Biol Crystallogr 67, 235–242.

Zhang, K. (2016). Gctf: Real-time CTF determination and correction. J Struct Biol 193, 1–12.

Zheng, S. Q., Palovcak, E., Armache, J. P., Verba, K. A., Cheng, Y., and Agard, D. A. (2017). MotionCor2: anisotropic correction of beam-induced motion for improved cryo-electron microscopy. Nat Methods 14, 331–332.

Zuckerman, D. M., Boucher, L. E., Xie, K., Engelhardt, H., Bosch, J., and Hoiczyk,E. (2015). The bactofilin cytoskeleton protein BacM of Myxococcus xanthus forms an extended β-sheet structure likely mediated by hydrophobic interactions. PLoS One 10, e0121074.

