## Supplementary Information for "Bactofilins form non-polar filaments that bind to membranes directly"

**Contents:**

Supplementary Figure S1-S6
Supplementary Tables T1-T2
Supplementary Movies M1-M4
Supplementary Data D1-D2
Supplementary References

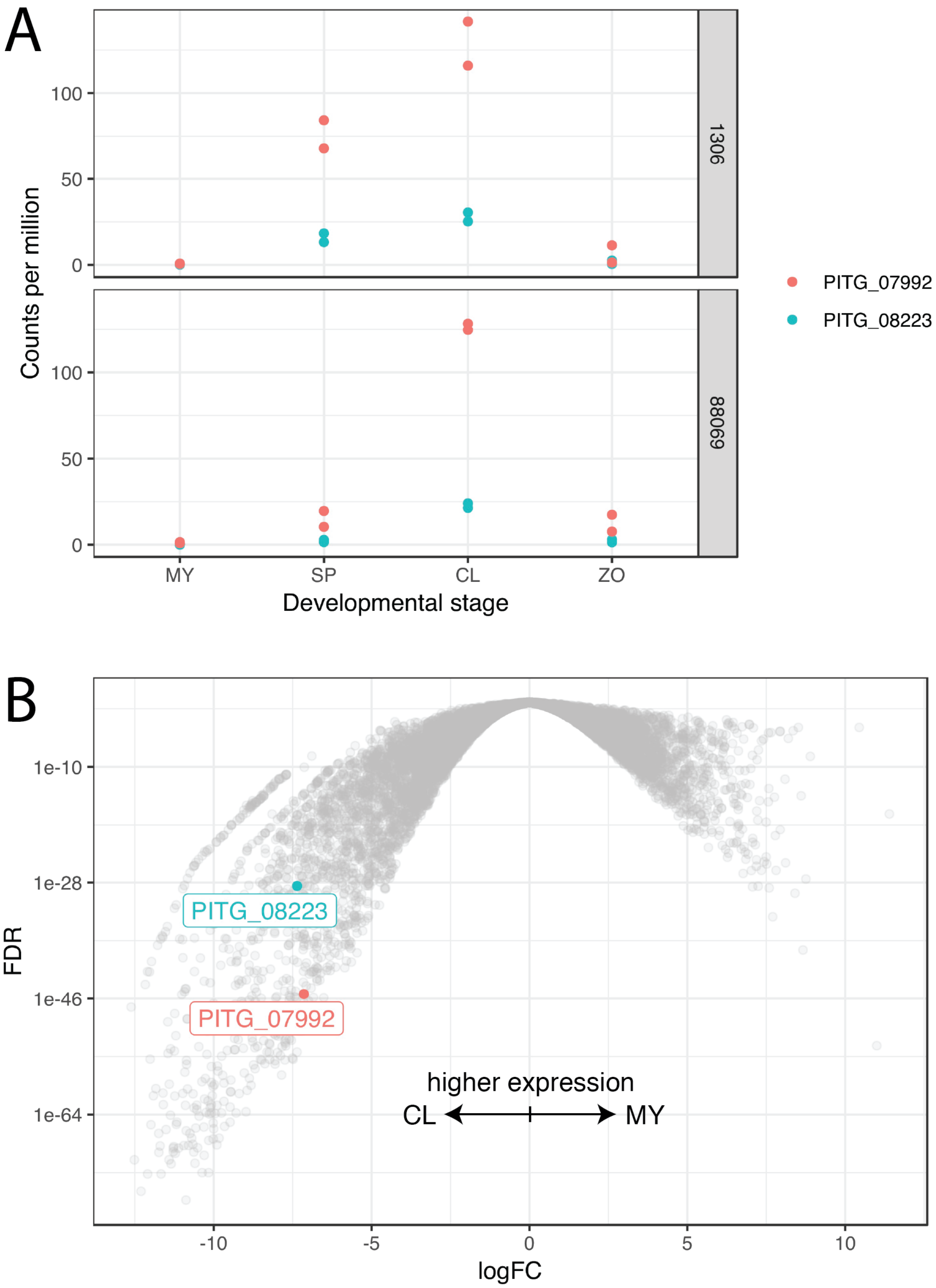


**Supplementary Figure S1.** A) Developmental changes in relative abundance (counts per million) of bactofilin mRNA. Data from (Ah-Fong et al., 2017). Upper and lower panels (datasets 1306 and 88069) correspond to independent experiments. Development stages: non-sporulating mycelia (MY), purified sporangia (SP), sporangia chilled in water to induce the cleavage of sporangia into zoospores (CL), zoospores released from the sporangia (ZO), and germinated cysts (GC). B) Volcano plot showing differential expression between MY and CL life stages in dataset 88069, data taken again from (Ah-Fong et al., 2017). Bactofilin genes are highlighted by colour and labels. Fold changes in expression of both bactofilin genes are in the 5$^{th}$ percentile of all genes.

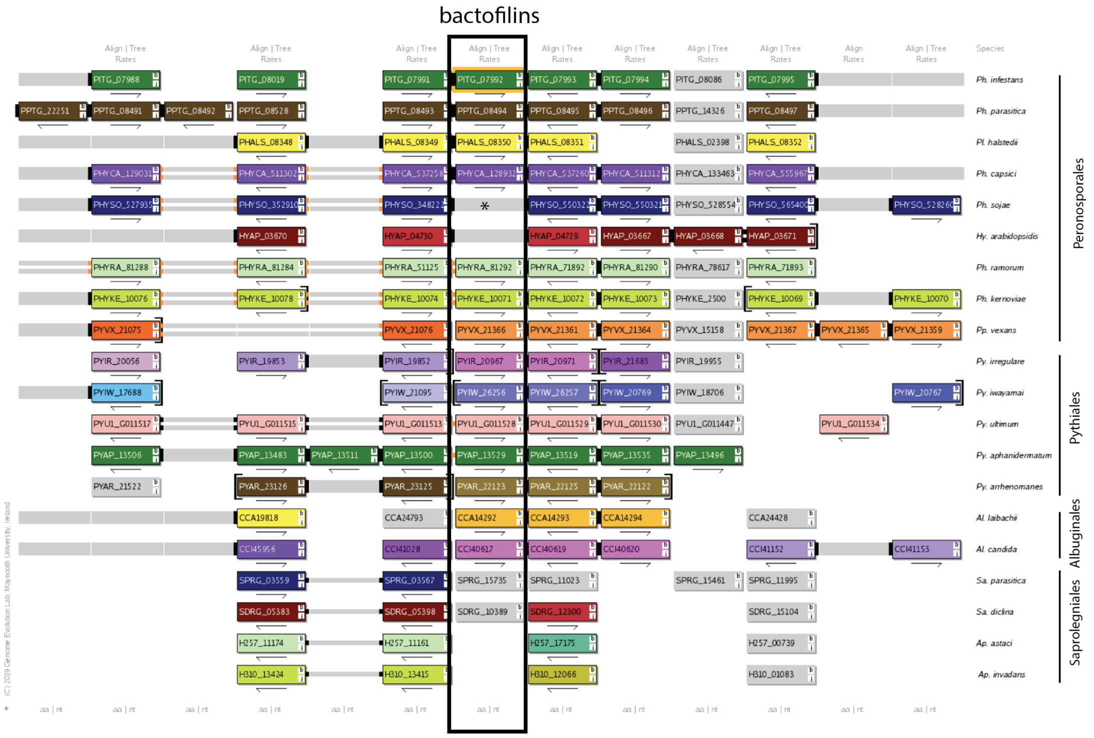


**Gene Box**
Genes are represented by coloured boxes that show its gene ID. Horizontal tracks correspond to chromosome / scaffold segments. Each genome has a colour palette that distinguishes different chromosomes or scaffolds. Genes that are in the same vertical column (homology pillar) are orthologs. Genes that are coloured grey are not syntenic.

**Connectors**
Connectors join nearby genes. A solid connector between two genes indicates that the two genes are adjacent in the genome, two smalls bars indicate that they are within 5 genes of each other and one small bar if they are within 20 genes. Connectors that are coloured orange denote an inversion. The connectors are continued over any intervening space between genes with grey extensions. Arrows: The arrows under each gene box denote the gene's relative orientation (i.e. Watson or Crick strand). Brackets: A bracket around a gene box indicates the end of a chromosome or scaffold.

From https://ogob.ie/gob/OGOB_help.pdf

**Supplementary Figure S2.** Output from the Oomycete Genome Order Browser (McGowan et al., 2019) for PITG_07992 (Uniprot D0N980). Orthologoues are found and at least some synteny of the region is present across the Perenosporales (with the exception of the non-spore forming *H. Arabidopsis*, which does not have a bactofilin), Pythiales and Albuginales. Orthologues are also found in Saprolegniales, but gene order is not conserved (Aphanomyces orthologues not shown in this view, and SDRG_10389 should be SDRG_10390). A star indicates the annotation in the *P. sojae* genome of PHYSO_348222 as a gene fusion, the downstream part is a bactofilin domain.

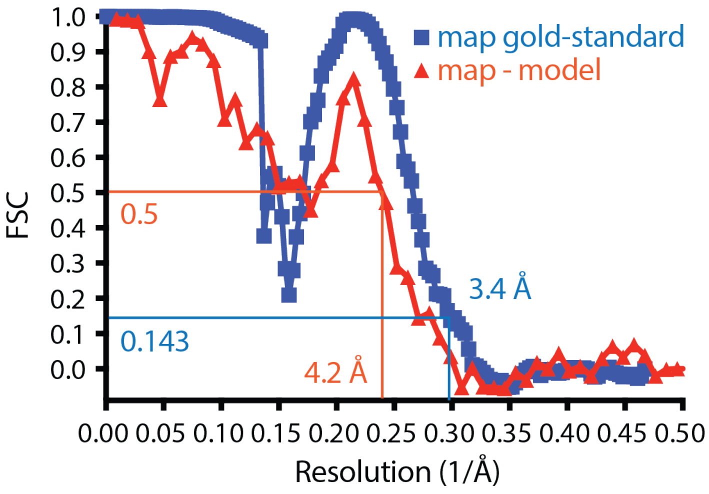


**Supplementary Figure S3.** Fourier Shell Correlation (FSC) analysis of the final cryo-EM TtBac bactofilin map and model. FSC 0.143 (Rosenthal and Henderson, 2003) leads to a nominal resolution of the map of 3.4 Å, but comparison of the map to the model indicates lower resolution around 4.2 Å, most likely caused by severe anisotropy of the cryo-EM map due to the beta stacking along the filament. This can be seen by the dip of the FSC curves before the peak at 4.7 Å, corresponding to the distance between the windings of the beta helix of bactofilin that dominates the Fourier transform of the filaments.

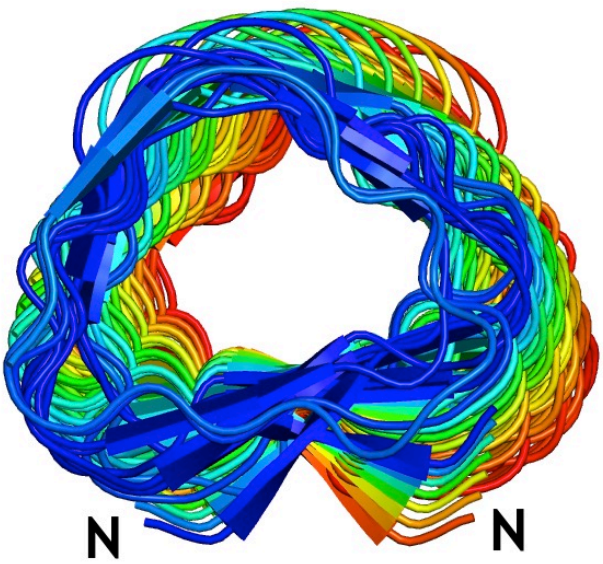


**Supplementary Figure S4.** In our cryo-EM structure, TtBac-WT protofilaments twist only very slightly, roughly 5° per 57 Å rise (see also Supplementary Table T1 for exact values, the rise corresponding to two bactofilin subunits because of the antiparallel arrangement of the subunits).

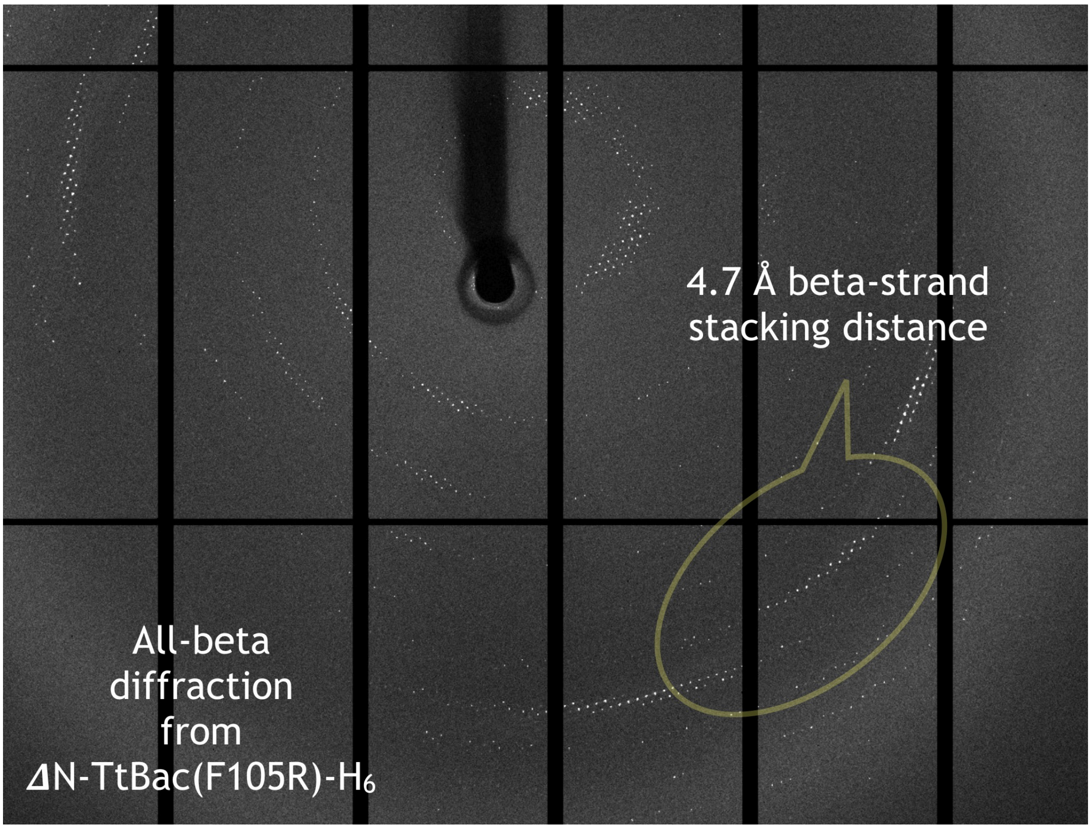


**Supplementary Figure S5.** All-beta diffraction from a bactofilin crystal similar to the ones used for structure determination (Supplementary Table T1). Diffraction is anisotropic and strongest around 4.7 Å, caused by the dominant beta-stacking interactions along the beta helical fold.

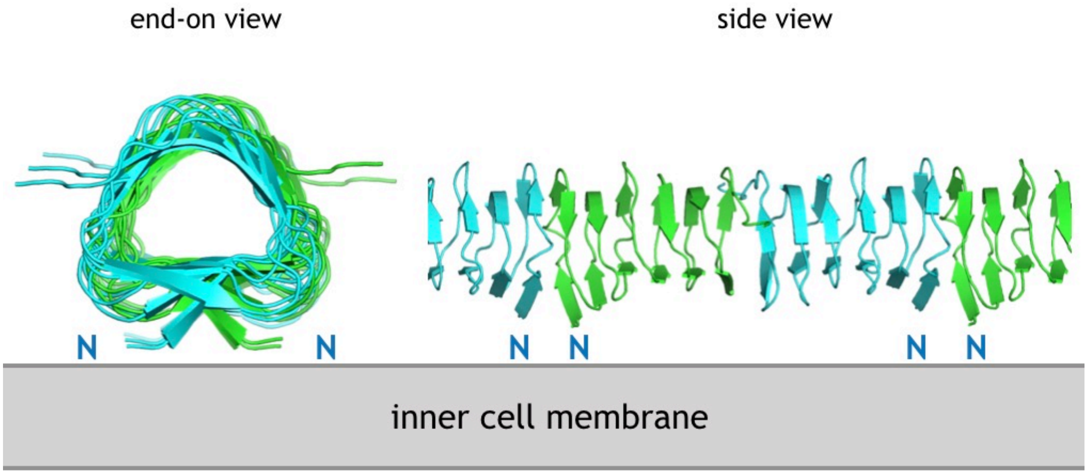


**Supplementary Figure S6.** Model of membrane binding by bactofilin filaments. The N-terminal tails containing a conserved membrane-targeting sequence (Figure 6A) bind to lipid membrane, utilising avidity that allows each interaction to be rather weak. The lack of significant twist of the filament (Supplementary Figure S4) allows the filament to bind over long distances.

### **Supplementary Table T1.** Cryo-EM and crystallography data

| **Statistics** | | | |
|---|---|---|---|
| **Sample** | ***Thermus thermophilus*** **bactofilin** TtBac | ***Thermus thermophilus*** **bactofilin** ΔN-TtBac(F105R)-$H_6$ SeMet | ***Thermus thermophilus*** **bactofilin** ΔN-TtBac(F105R)-$H_6$ native |
| NCBI database ID | WP_011173792.1 | WP_011173792.1 | WP_011173792.1 |
| Constructs | 1-123 | 11-123-GSHHHHHH, F105R mutation | 11-123-GSHHHHHH, F105R mutation |
| **Method** | cryo-EM | crystallography SeMet SAD | crystallography native |
| **Data collection** | | | |
| Beamline/microscope | FEI Krios, Falcon III ec | Diamond I03 | Diamond I03 |
| Wavelength / energy | 300 kV | 0.97928 Å<br>3 crystals merged | 0.97623 Å |
| **Crystal / helical** | | | |
| Space / point group | RELION D1 symmetry | $I2_12_12_1$ | $I2_12_12_1$ |
| Cell (Å) | (C2 along X) | 198.0, 247.3, 504.5 | 191.9, 244.9, 505.9 |
| Twist / rise | 4.89°, 57.46 Å | | |
| **Data** | | | |
| Resolution (Å) | 3.4 map / 4.2 model | 4.0 | 3.5 |
| Completeness (%)[1] | | 99.9 (100.0) | 99.2 (99.2) |
| Multiplicity[1] | | 40.4 (40.7) | 3.8 (3.9) |
| (I) / σ(I) [1] | | 10.8 (2.9) | 7.1 (2.5) |
| $R_{merge}$[1] | | 0.418 (2.086) | 0.123 (0.541) |
| $R_{pim}$[1] | | 0.094 (0.469) | 0.070 (0.309) |
| CC1/2 | | 0.998 (0.901) | 0.996 (0.869) |
| Anomalous correlation | | 0.037 (-0.008) | |
| Selenium sites | | 32 | |
| Images, pixel size | 2130, 1.07 Å | | |
| Defocus range, dose | -1.0 - -3 μm, ~40 e/Å$^2$ | | |
| Helical segments | 346k, 57 Å apart | | |
| **Refinement** | | | |
| R / $R_{free}$[2] | real-space refined | | 0.283 / 0.307 |
| Models | 2 chains, 12-112, no waters | | 32 chains, aa 11-100, no waters |
| Bond length rmsd (Å) | 0.007 | | 0.003 |
| Bond angle rmsd (º) | 1.041 | | 0.682 |
| Favoured (%)[3] | 100.0 | | 100.0 |
| Disallowed (%)[3] | 0.0 | | 0.0 |
| MOLPROBITY | 96th percentile | | 100th percentile |
| **PDB/EMDB IDs** | **6RIB, EMD-4887** | | **6RIA** |

[1] Values in parentheses refer to the highest recorded resolution shell.
[2] 5% of reflections were randomly selected before refinement.
[3] Percentage of residues in the Ramachandran plot (PROCHECK 'most favoured' and 'additionally allowed' added together).

**Supplementary Table T2. Proteins used in this study**

PiBac: *Phytophthora infestans bactofilin*, NCBI ref: EEY54368.1
*TtBac: Thermus thermophilus* bactofilin, NCBI ref: WP_011173792.1

**PiBac-WT 1-203 negative stain EM**
MEEAPVPRNPPPKPKRSNVPSAPADYPDDTYSDQDYNMSPIRRGRQHANR
QSGSPPMTPPYTVPHQAKVPIIDAEPETTIGAAVKMKGELSFERLLRIEG
EFEGKLNSKGSLVIGTRGALIGNVDNMKEVYITGGRIVGNVNVEKLVLRD
KAQIFGNIIAKSVKIEPECIVVGRINVNPQAPERINEKGEIVKDDAPDGT
PSS

**$H_6$-TtBac 1-123 negative stain EM, cryo-EM**
MGSSHHHHHHMGRMLGRKERTLTYLGPDTEVLGDMRAKGQVRIDGLVRGS
VLVEGELEVGPTGRVEGERVEARSVLIHGEVKAELTAEKVVLSKTARFTG
QLKAQALEVEAGAVFVGQSVAGEHKALEAPKEA

**TtBac-WT 1-123 cryo-EM, membrane binding studies**
MGRMLGRKERTLTYLGPDTEVLGDMRAKGQVRIDGLVRGSVLVEGELEVG
PTGRVEGERVEARSVLIHGEVKAELTAEKVVLSKTARFTGQLKAQALEVE
AGAVFVGQSVAGEHKALEAPKEA

**Nanobody NB4-mut2-(L13S, Q15D, K45D, K66D) cryo-EM**
MAQVQLQESGGGSVDAGGSLRLSCAASGRTFGASLMGWFRQAPGDEREFV
AAINWTGKIWYTDSVDGRFTISRDNAKNTANLQMNNLTPEDTAIYYCAAR
LGIGFAPSSVEYDYWGQGTQVTVSSAAASSHHHHHH

**ΔN-TtBac(F105R)-$H_6$ 11-123 crystallography, polymerisation-impaired**
MTLTYLGPDTEVLGDMRAKGQVRIDGLVRGSVLVEGELEVGPTGRVEGER
VEARSVLIHGEVKAELTAEKVVLSKTARFTGQLKAQALEVEAGAVRVGQS
VAGEHKALEAPKEAGSHHHHHH

**ΔN-TtBac 11-123 membrane binding studies**
MTLTYLGPDTEVLGDMRAKGQVRIDGLVRGSVLVEGELEVGPTGRVEGER
VEARSVLIHGEVKAELTAEKVVLSKTARFTGQLKAQALEVEAGAVFVGQS
VAGEHKALEAPKEA

**Supplementary Movies:**

**Supplementary Movie M1.** Cryo-EM density after helical reconstruction of TtBac filament bound by nanobody NB4-mut2. Boundaries of the bactofilin monomers are clearly visible, as is their antiparallel arrangement in each protofilament. An image from this movie is shown in Figure 3F.

**Supplementary Movie M2.** Overview of TtBac bactofilin cryo-EM structure, providing a better 3D impression of the assembly.

**Supplementary Movie M3.** Electron cryotomography (cryo-ET) of an *E. coli* cell with TtBac-WT overexpressed. Note that filament bundles are arranged all around the cell's periphery, under the inner membrane and are particularly obvious when the movie goes through the upper and lower cellular envelope where the bactofilin bundles run at roughly 45° angles to the long cell axis. Images from the tomogram are shown in Figure 6E, left.

**Supplementary Movie M4.** Same as Movie M3, but ΔN-TtBac has been overexpressed. Because the filaments no longer bind to the inner membrane of the *E. coli* cells, a very large bactofilin bundle runs along the long cell axis, also inhibiting cell division at the septum site. An image from this tomogram is shown in Figure 6E, right.

**Supplementary Data:**

**Supplementary Data D1. Gene tree of all bactofilins.** Gene tree showing inferred phylogeny of a representative set of bacterial and archaeal bactofilins, and all identified putative eukaryotic bactofilins. Tips are labelled: "<NCBI taxid> | <NCBI Species> | <Gene accession/uniprot identifier>". Tips corresponding to non-bacterial sequences are marked with coloured circles, coloured by NCBI taxonomy level 4.

**Supplementary Data D2. Eukaryotic bactofilins.** CSV (comma separated values) file of putative eukaryotic bactofilins.

### Supplementary References

Ah-Fong, A. M., Kim, K. S., and Judelson, H. S. (2017). RNA-seq of life stages of the oomycete Phytophthora infestans reveals dynamic changes in metabolic, signal transduction, and pathogenesis genes and a major role for calcium signaling in development. BMC Genomics *18*, 198.

McGowan, J., Byrne, K. P., and Fitzpatrick, D. A. (2019). Comparative Analysis of Oomycete Genome Evolution Using the Oomycete Gene Order Browser (OGOB). Genome Biol Evol *11*, 189-206.

Rosenthal, P. B., and Henderson, R. (2003). Optimal determination of particle orientation, absolute hand, and contrast loss in single-particle electron cryomicroscopy. J Mol Biol *333*, 721-745.
