## Supplementary figures and images for "Bactofilins form non-polar filaments that bind to membranes directly"

### Supplementary Data 1

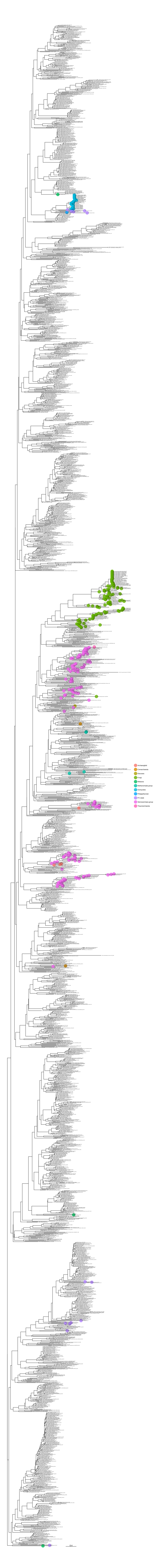
